# CoVizu: Rapid analysis and visualization of the global diversity of SARS-CoV-2 genomes

**DOI:** 10.1101/2021.07.20.453079

**Authors:** Roux-Cil Ferreira, Emmanuel Wong, Gopi Gugan, Kaitlyn Wade, Molly Liu, Laura Muñoz Baena, Connor Chato, Bonnie Lu, Abayomi S. Olabode, Art F. Y. Poon

## Abstract

Phylogenetics has played a pivotal role in the genomic epidemiology of SARS-CoV-2, such as tracking the emergence and global spread of variants, and scientific communication. However, the rapid accumulation of genomic data from around the world — with over two million genomes currently available in the GISAID database — is testing the limits of standard phylogenetic methods. Here, we describe a new approach to rapidly analyze and visualize large numbers of SARS-CoV-2 genomes. Using Python, genomes are filtered for problematic sites, incomplete coverage, and excessive divergence from a strict molecular clock. All differences from the reference genome, including indels, are extracted using minimap2, and compactly stored as a set of features for each genome. For each Pango lineage (https://cov-lineages.org), we collapse genomes with identical features into ‘variants’, generate 100 bootstrap samples of the feature set union to generate weights, and compute the symmetric differences between the weighted feature sets for every pair of variants. The resulting distance matrices are used to generate neigihbor-joining trees in RapidNJ and converted into a majority-rule consensus tree for the lineage. Branches with support values below 50% or mean lengths below 0.5 differences are collapsed, and tip labels on affected branches are mapped to internal nodes as directly-sampled ancestral variants. Currently, we process about million genomes in approximately nine hours on 34 cores. The resulting trees are visualized using the JavaScript framework D3.js as ‘beadplots’, in which variants are represented by horizontal line segments, annotated with beads representing samples by collection date. Variants are linked by vertical edges to represent branches in the consensus tree. These visualizations are published at https://filogeneti.ca/CoVizu. All source code was released under an MIT license at https://github.com/PoonLab/covizu.

## INTRODUCTION

Severe acute respiratory syndrome coronavirus 2 (SARS-CoV-2) was first sampled in December, 2019, in association with an outbreak of unexplained pneumonia in the province of Hubei, China [1]. The first genome sequence of the novel coronavirus isolated from this outbreak was released into the public domain on January 10, 2020 [2]. Early phylogenetic analyses of this and subsequent genome samples provided initial evidence of human-to-human transmission [3] and estimates of the basic reproduction number [4]. One of the most remarkable developments from the global pandemic has been the rapid accumulation and generally timely release of SARS-CoV-2 genome sequence data into the public domain. As of June 21, 2021, over two million SARS-CoV-2 genome sequences have been deposited in the Global Initiative on Sharing All Influenza Data (GISAID) database [5], and this number has grown at a sustained and exponentially increasing rate (Figure 1). The phylogenetic analysis of these data has played an important role in tracking the genomic epidemiology of SARS-CoV-2. For example, Nextstrain (https://nextstrain.org/) [6] publishes time-scaled phylogenetic trees [7] as interactive visual summaries of the diversity of SARS-CoV-2 genomes at global and local scales. The resulting web documents are updated in real time with the availability of new data. Throughout the SARS-CoV-2 pandemic, Nextstrain has featured prominently in global variant tracking and scientific communication, and in some cases it has directly influenced public health decision-making [8].

**Figure 1:**
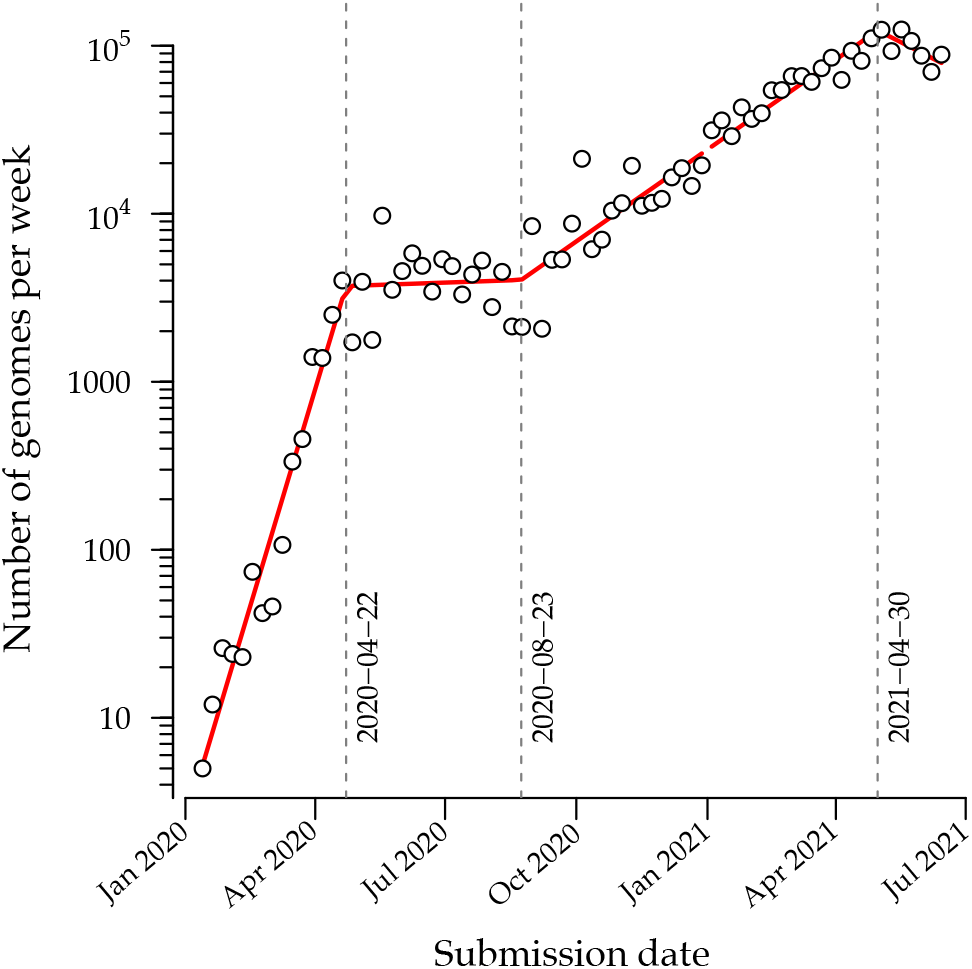
Weekly numbers of genomes submitted to the GISAID database (accessed on 2021-06-26). Red line segments represent the fit of a piecewise linear regression with three change points (indicated by vertical dashed lines) using the R package segmented [9]. An increasing linear trend relative to a log-transformed y-axis indicates an exponentially growing rate of genome submission.

However, the data visualization maintained by Nextstrain is limited to about 3,000 genome sequences due to the computational complexity of calculating a maximum likelihood tree, and the constraints of drawing large trees in a limited visual space. Accurately reconstructing a large phylogeny is difficult because the number of possible trees grows faster than exponentially with the number of observed sequences; however, the amount of phylogenetic signal in the data has a fixed upper limit since we cannot sequence more than the full length genome. Furthermore, the time scale of transmission for SARS-CoV-2 tends to outpace its molecular clock, such that many new infections are genetically identical to their source populations. Paradoxically, we can become increasingly uncertain about the relationships among specific lineages as we collect greater amounts of data [10]. This uncertainty is exacerbated by sequencing error [11] and a substantial prevalence of missing data (incomplete genome sequences).

Even if it is computationally feasible to accurately infer the evolutionary relationships among millions of sampled infections, visualizing these results in a meaningful way is a significant challenge. A standard maximum likelihood phylogeny, for example, does not differentiate between samples with identical sequences. These samples become collapsed into a single node, even if they were collected on different dates or at different locations. Given these meta-data, however, genetically identical samples carry epidemiologically relevant information at an individual level. While Bayesian methods can incorporate prior information to resolve the relationships between identical samples [12], these methods are computationally demanding and the outputs do not necessarily result in an efficient nor effective use of visual space.

Here, we describe an ongoing open-source project to provide a public interface to visualize the global diversity of SARS-CoV-2 genomes in near real time. Development of CoVizu (derived from ‘coronavirus visualization’) began in April 2020. From December 2020 onwards, CoVizu became provisioned by a customized data feed from the GISAID database (https://gisaid.org), which is presently the largest publicly-accessible repository of SARS-CoV-2 genome sequence data in the world. The specific objectives of this project are: (1) to process and visualize as much of the GISAID database as possible (*i.e.*, millions of genomes); (2) to reconstruct robust evolutionary and epidemiological relationships among these genomes; (3) to continually update outputs with new genomic data as frequently as possible, and; (4) to present this information in a rich and intuitive visual interface.

CoVizu is composed of a Python-based ‘backend’ — an analytical pipeline for rapidly inferring the evolutionary relationships among genome sequences — and a JavaScript-based ‘frontend’ to visualize these relationships. It relies heavily on the manually curated Pango nomenclature system that partitions the global diversity of SARS-CoV-2 into a hierarchy of ‘lineages’ [13]. The web interface is presently hosted at https://filogeneti.ca/CoVizu, and as an integrated component of the GISAID web portal. All Python and JavaScript source code comprising the project is publicly available from our repository (https://github.com/PoonLab/CoVizu) under the MIT license.

## DATA ANALYSIS

The CoVizu backend is implemented in the Python scripting language. Raw sequence data and metadata, including sample collection dates and Pangolin [13] lineage assignments, are provisioned by the GISAID database as a single Lempel-Ziv-Markov compressed JSON (JavaScript Object Notation) file.

### Sequence alignment and cleaning

An uncompressed data stream from the GISAID provisioned file is processed in Python to exclude any record whose genome sequence: (1) lacks a Pango lineage assignment; (2) was sampled from a non-human host; (3) was shorter than 29,000 nt; (4) lacks a complete sample collection date (*e.g.*, year and month with no day); (5) or was labeled with a collection date preceding 2019-12-01 or in the future. The filtered data stream is then redirected to the program minimap2 (version 2.17) [14] for pairwise alignment against the SARS-CoV-2 reference genome (Genbank accession NC_045512 [2]). We parse the resulting SAM (sequence alignment/map) formatted output stream in Python to extract any genetic differences (nucleotide substitutions, insertions and deletions) from the reference as ‘features’, as well as any intervals of uncalled bases (missing data). These provide a compact representation of each genome sequence. Any genomes that failed to map to the reference or contained over 300 (∼1%) uncalled bases are excluded at this stage. Furthermore, the feature set of each genome is screened for problematic sites using the curated list maintained at https://github.com/W-L/ProblematicSites_SARS-CoV2 [15]. We also exclude genomes where the number of features exceeds the 99.5% percentile of a Poisson distribution with rate parameter *λ* = *r*Δ*t*, where *r* = 0.0655 substitutions/genome/day and Δ*t* is the number of days since 2019-12-01 [16]. We use the SciPy root-finding method [17] to numerically solve for the transition points of the Poisson cumulative distribution function at different values of Δ*t*. The genome records that pass these filters are partitioned into a dictionary keyed by Pango lineage assignment. This dictionary is also serialized to a JSON file for parallel computation with a message passing interface (MPI).

### Time-scaled tree reconstruction

To reconstruct a time-scaled phylogeny that relates the Pango lineages in the database, we select for each lineage the earliest sample passing all the above filtering criteria as the representative genome of each lineage. We reconstitute a multiple sequence alignment of these representative genomes from the respective feature sets by excluding all insertions relative to the reference genome. Next, we reconstruct a maximum likelihood tree using the fast approximate heuristic method implemented in FastTree (version 2.1.11, compiled for double precision) [18]. We rescale the resulting tree using TreeTime (version 0.8.0) [7] with a pre-specified clock rate (8 × 10^−4^ substitutions/site/year). This final tree is processed using the Biopython Phylo submodule [19] and serialized into the Newick tree format with terminal nodes labeled by Pango lineage.

### Clustering analysis

We use the neighbor joining method [20] to reconstruct the evolutionary relationships among genomes within each Pango lineage. This clustering method requires a pairwise distance matrix. Because we want to incorporate indel variation while minimizing computation time, we assume a uniform rate over all possible genetic differences and ignore multiple hits. To minimize the memory consumption at this step, we convert the features into integers by indexing the ordered set union of all features observed for a given lineage. All genomes with identical feature sets are compressed into a single “variant”. For each lineage, we compute the symmetric difference (*A* Δ *B*) between the feature sets for every pair of variants, where *A* Δ *B* contains all features that are in either *A* or *B* but not both. For example, the symmetric difference between the subsets *A* = {1, 4, 128} and *B* = {4, 37, 89} is *A* Δ *B* = {1, 37, 89, 128}. To generate a bootstrap replicate, we sample the feature set union at random with replacement, and use the resulting frequencies to weight the symmetric differences. Thus, the distance between *A* and *B* is given by Σ_*i*∈*A*Δ*B*_ *f_b_*(*i*), where *f_b_*(*i*) is the frequency of the *i*-th feature in bootstrap replicate *b*. The resulting pairwise distance matrix is written to a comma-delimited file as input for neighbor-joining tree reconstruction using RapidNJ (version 2.3.2) [21]. We repeat this process for 100 bootstrap replicates. To reduce computing time, all lineages with 5,000 samples or fewer are processed in a single batch distributed over multiple cores in a clustered computing environment using the Python-MPI bindings implemented in the mpi4py module [22]. Lineages with more than 5,000 samples are processed singly, with bootstrap replicates distributed over multiple cores.

We use a custom Python function to generate a consensus tree from all splits that occurred in at least 50% of the bootstrap trees, and assign branch lengths by averaging over the subset of trees containing each split. Next, we collapse any branches with a mean length below 0.5 features (genetic differences). If a terminal branch is collapsed, then its variant label is re-assigned to the parent internal node. If an internal branch is collapsed, then any variant labels carried by that node are re-assigned to its parent. Thus, an internal node may be associated with multiple variants. We interpret a labeled internal node as an ancestral variant that has been directly observed as a genome sequence. The resulting tree is serialized into a JSON file comprising node and edge lists. A node list is an associative array comprising lists of sample labels keyed by variant. An edge list comprises pairs of parent and child nodes (variants), branch lengths and bootstrap support values.

## DATA VISUALIZATION

The CoVizu front-end is implemented in JavaScript using the D3.js (https://d3js.org/) and jQuery UI (https://jqueryui.com/) frameworks. Upon completion of the analysis pipeline, JSON data from the clustering analysis and the Newick file from the time-scaled tree reconstruction are automatically transferred from the computing cluster to the webserver. To reduce page load time, the JSON data is transmitted to the client in a gzip compressed format. These data are used to render SVG (support vector graphics) and HTML elements in three panels that represent different levels of data aggregation from left to right. The leftmost panel depicts the time-scaled tree relating Pango lineages and correspond to the highest level of data aggregation. The middle panel depicts a ‘beadplot’ that we use to visualize the genetic variation within a selected Pango lineage. The width of the beadplot scales dynamically with the horizontal dimension of the browser window. Finally, the rightmost panel displays an interactive dynamic table that displays the individual samples for the selected lineage, variant, or bead. All these visual outputs are presented as a single composite web page (Figure 2). This webpage and its text components, *e.g.*, pop-up dialogs, have been translated into French, Spanish, and Chinese.

**Figure 2:**
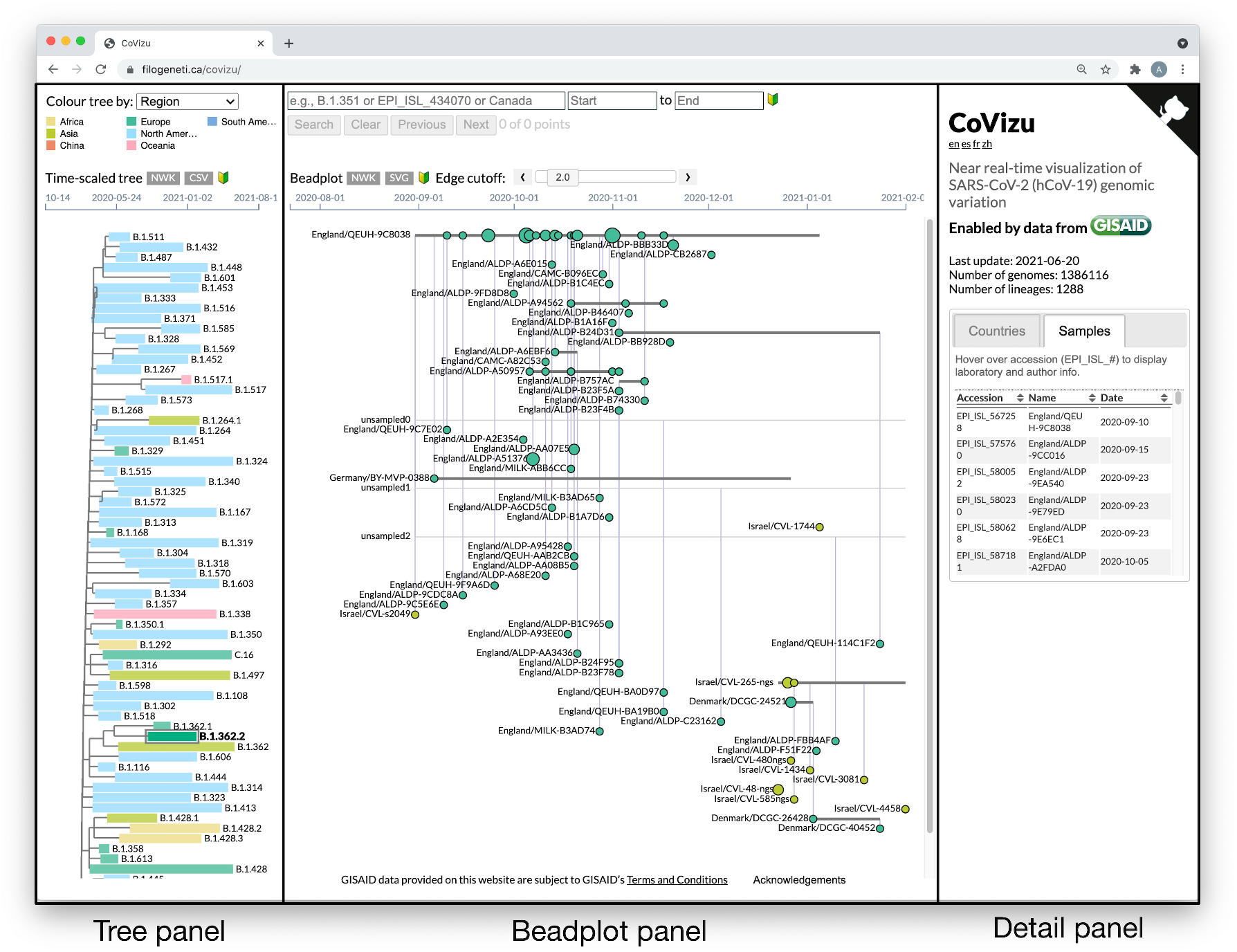
The CoVizu front-end presented as a single webpage at https://filogeneti.ca/CoVizu. Visual information is arranged into three panels (emphasized with rectangular boxes and labels) to present the data at decreasing levels of granularity from left to right. The leftmost panel displays a time-scaled tree relating Pango lineages, coloured by geographic region in this instance. Selecting a lineage updates the middle panel to display a beadplot of its variants and samples. In this example, we have selected lineage B.1.362.2, which was sampled predominantly in Europe and comprised 99 samples grouped into 59 variants. The rightmost panel depicted here displays a scrollable table of sample accessions, names and collection dates.

### Time-scaled tree

The time-scaled tree relating Pango lineages is rendered as an SVG using a rectangular layout algorithm, with the earliest time point on the lefthand side. Each tip representing a lineage is associated with a rectangular element spanning the range of sample collection dates. The user can select whether the rectangles are coloured according to the number of samples, most recent collection date, average deviation from the molecular clock, or the predominant geographical region of sampling (Africa, Asia, China, Europe, North America, Oceania, and South America). China is classified as a region because the samples from this country in the GISAID database are labelled by district (*e.g.*, Guangzhou) instead of by country. We used a qualitative colour-accessible palette developed by Paul Tol (https://personal.sron.nl/~pault/) for regions, and built-in D3.js palettes for other colour schemes. The time-scaled tree can be downloaded by the user in the Newick tree format, and lineage-level statistics can be downloaded as a comma-separated values (CSV) formatted file.

Mouseover events on rectangular elements trigger a ‘tool tip’ dialog that provides lineage-level summary statistics, such as the number of samples and mean deviation from the clock model. In addition, this dialog displays a list of all mutations that were observed in at least 50% of samples. Following the colon-delimited notation used in https://cov-lineages.org, amino acid substitutions are prefixed with ‘aa’ and labeled by the protein abbreviation and position in the reference protein sequence. For example, ‘aa:S:D614G’ represents a substitution of aspartic acid by glycine at position 614 of the spike protein. Insertions and deletions are prefixed respectively with ‘ins’ and ‘del’, and labeled by the reference nucleotide coordinate and indel length in nucleotides. For example, ‘del:11288:9’ represents a deletion of 9 nucleotides at genome coordinates 11288-11296 (inclusive).

### Beadplots

Selecting a lineage in the time-scaled tree triggers the browser to render a beadplot (Figure 3) in the middle panel as an SVG. A beadplot is a custom visual device that summarizes the distribution and genetic variation of samples within a lineage. The horizontal axis of the beadplot is scaled to the range of sample collection dates for the lineage. Samples with identical genome sequences are grouped into variants. In other words, each variant corresponds to a node in the consensus tree. (In the context of SARS-CoV-2, the term *variant* is often used interchangeably with *clade* or *lineage* [23]. However, *variant* can also refer to any unique combination of differences from a reference sequence.) Each variant is represented by a horizontal line segment in the beadplot.

**Figure 3:**
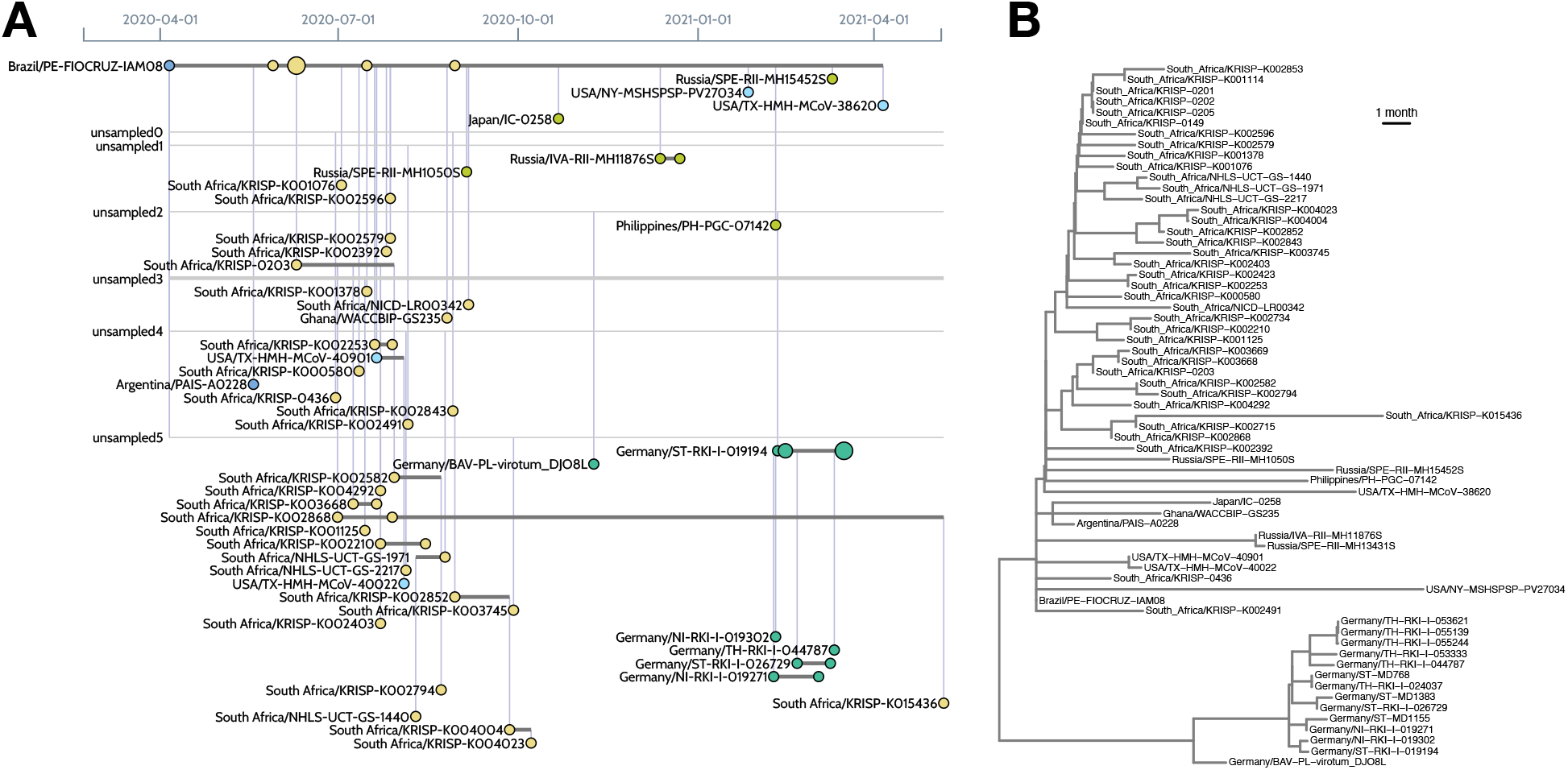
Visualizing the sample composition of a lineage. (A) The left image was generated by the CoVizu website for lineage B.1.1.117, using data retrieved from GISAID on June 20, 2021. Each horizontal line represents genomes that are indistinguishable in sequence (comprising a ‘variant’), labeled by the name of the earliest sample. For example, Brazil/PE-FIOCRUZ-IAM08 (upper left) was sampled on 2020-04-06, and identical genomes were subsequently observed in 6 samples in South Africa. This pattern is consistent with the importation of this variant from Brazil to South Africa. A more recent variant (Germany/ST-RKI-I-019194, 7 samples) is ancestral to several other variants sampled in Germany (lower right). It is derived from an unsampled ancestral variant (‘unsampled2’) at a distance of 3.8 mutations, averaged over 100 bootstrap replicates, which in turn is separated from Brazil/PE-FIOCRUZ-IAM08 by a mean of 10.5 mutations. These lengths imply that this lineage is relatively undersampled. (B) For comparison, the right image depicts a time-scaled tree generated from the same data using FastTree2 and TreeTime. In this visualization, it is more difficult to identify samples that are genetically indistinguishable. Thus, beadplots endeavour to visually emphasize features that are relevant for public health applications.

Each horizontal line segment spans the range of sample collection dates. Circles (beads) along a line segment represent samples. The area of the bead is scaled in proportion to the number of samples collected on the same date. In addition, each circle is coloured with respect to the most common geographic region of the samples. Together these elements provides an intuitive visual summary of the frequencies of a specific variant over time (Figure 3A). Furthermore, sampling the same or closely related variants in different regions provides evidence of importation events; for example, see Figure 3B. Unsampled variants, which correspond to unlabelled internal nodes in the consensus tree, are represented by horizontal line segments without beads. The existence of these latent variants is inferred by the common ancestry of variants that are directly observed. Variants are connected by vertical line segments that correspond to branches in the consensus tree. Because the number of branches can become excessive for large beadplots, we provide a slider widget for users to filter branches by mean length. Horizontal lines can extend beyond the first and last sample of a variant if descendant variants are sampled at earlier or later dates, respectively.

All elements of a beadplot SVG are visually responsive to mouseover events, which also triggers a tooltip dialog summarizing variant- or bead-level summary statistics, including the number of samples, branch lengths, and the parent and child variants. The displayed beadplot can be exported as either a consensus tree (Newick format) or a SVG file, that can be converted into any rasterized or vector-based image format.

### Sample details

The rightmost panel of the web page presents some database-level statistics — namely, the date of the current update, total number of genomes that past the quality filters, and the number of Pango lineages — and a tabbed content area where the user can switch between two interfaces that summarize the sample metadata. The ‘countries’ interface displays a barplot that summarizes the distribution of samples among geographic regions, and a sortable table that breaks down these frequencies by country (Figure 4A) for the selected element. The ‘samples’ interface, on the other hand, displays a sortable table that lists the accession number, name (label) and collection date for every sample associated with the selected element. In addition, mouseover events are bound to the accession numbers in the table to trigger a query of the GISAID API for retrieving laboratory and author information for the sample, which is displayed as a pop-up dialog (Figure 4B).

**Figure 4:**
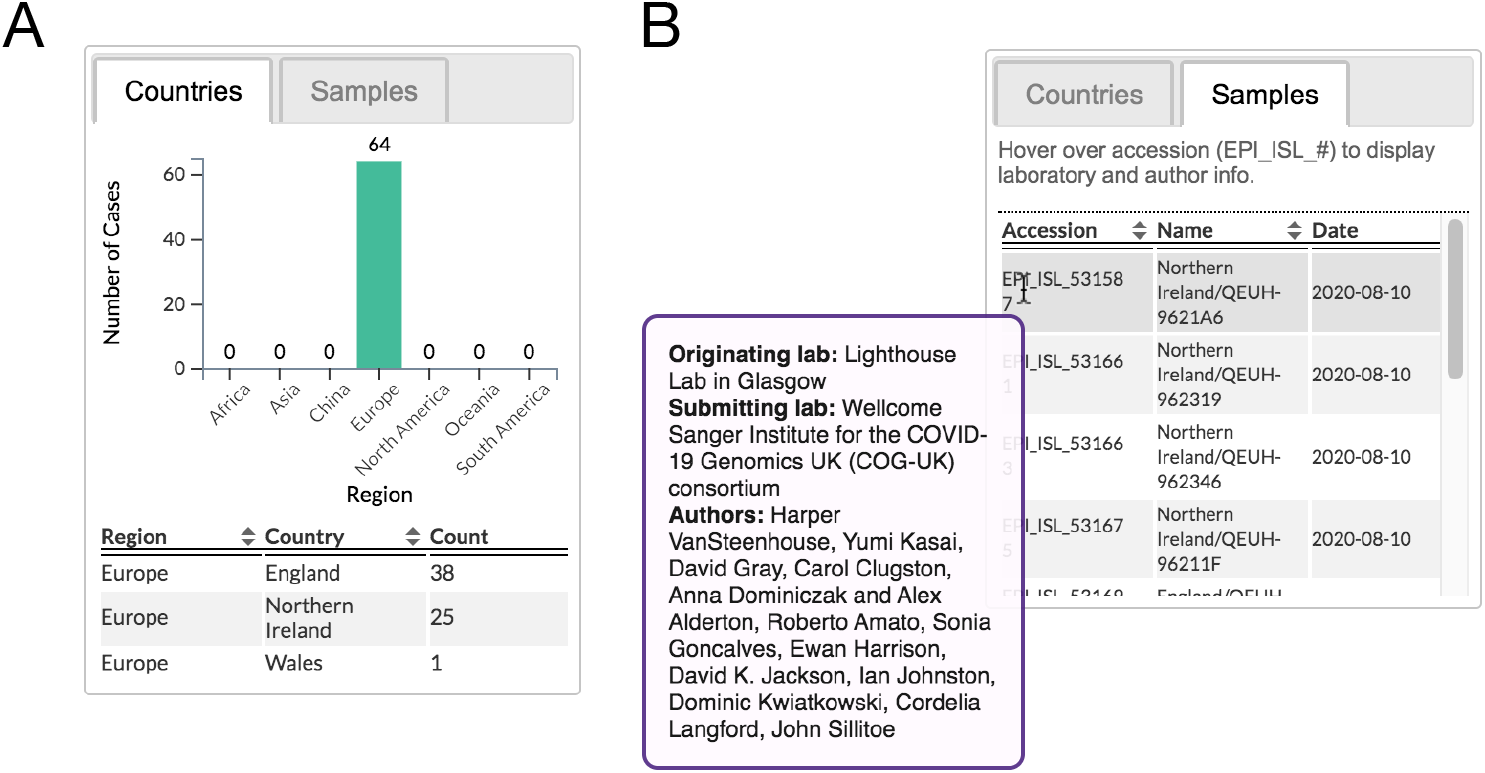
Detailed metadata display. (A) An example of the metedata displayed on the web-interface for a bead and (B) the contributing laboratory information provided for each sample give the accession number.

### Search interface

Since the front-end was designed to enable users to browse the relationships among millions of SARS-CoV-2 samples, we also needed to implement a search interface to enable users to quickly focus on samples matching specific parameters. The search interface comprises a text box for submitting a substring query, which can be matched against Pango lineage names, GISAID accession numbers, countries, and sample names; and date selection widgets for specifying a range of sample collection dates. If the substring query matches a regular expression that identifies it as a partial Pango lineage name or accession number, the browser populates a drop-down with suggested ‘autocompletions’ of the substring.

The submitted query is compared to metadata extracted from all samples, and the unique identifiers of bead and lineage elements that contain hits are stored. Next, the browser modifies the class attribute of all of matching elements, which causes the window to update how these elements are drawn, *i.e.*, with CSS highlighting. Caching the search results in this way streamlines the process of navigating between lineages and rendering the corresponding beadplots. The total number of hits is displayed below the search interface. Finally, the user can traverse search results using either the ‘next’ and ‘previous’ buttons or arrow keys.

## CONCLUDING REMARKS

Over the course of the SARS-CoV-2 pandemic, it has quickly become clear that the standard phylogenetic toolkit is not up to the task of processing the overwhelming number of publicly accessible viral genomes collected around the world. This critical situation has catalyzed the development of new analytical methods [12, 24]. It has also led to the resurrection of classic methods in phylogenetics, including maximum parsimony [25] and, in our case, neighbor joining with an uncorrected distance. CoVizu is under continual development and many of the methods described here are subject to refactoring for improved performance. We welcome suggestions for additional features, with the hope that this rapid analysis and visualization system can provide a unique, useful resource for public health monitoring and basic research.

## ACKNOWLEDGEMENTS

We gratefully acknowledge all the Authors, the Originating laboratories responsible for obtaining the specimens, and the Submitting laboratories for generating the genetic sequence and metadata and sharing via the GISAID Initiative, on which this research is based. Acknowledgement tables for the specific contributions to the GISAID database depicted in figures are provided as Supplementary Material. We also thank the GISAID technical development team for providing JavaScript code to retrieve laboratory and author information using their sample accession API. An earlier version of this work was presented at the 28th International Dynamics & Evolution of Human Viruses conference.

## AUTHOR CONTRIBUTIONS

All authors reviewed the manuscript. RF, EW and AFYP drafted the manuscript. AFYP conceived of the visualization scheme, designed the software, and wrote the initial backend and frontend source code. EW implemented Python scripts for database transactions and backend processing. GG and BL wrote unit tests for backend and frontend code. RC and LMB implemented and tested the search interface. LMB provided Spanish language translations. AO implemented genetic diversity analyses. KW implemented tooltips and contributed to the barplot and tabular interfaces. ML contributed CSS code and provided Chinese language translations. CC evaluated open-source implementations of neighbor-joining and ran validation experiments.

