## Supplemental Table 1 for "CoVizu: Rapid analysis and visualization of the global diversity of SARS-CoV-2 genomes"

We gratefully acknowledge the following Authors from the Originating laboratories responsible for obtaining the specimens, as well as the Submitting laboratories where the genome data were generated and shared via GISAID, on which this research is based.

All Submitters of data may be contacted directly via www.gisaid.org

Authors are sorted alphabetically.

| Accession ID | Originating Laboratory | Submitting Laboratory | Authors |
|---|---|---|---|
| EPI_ISL_1037838 | Laboratory Corporation of America | Respiratory Viruses Branch, Division of Viral Diseases, Centers for Disease Control and Prevention | Peter W. Cook, Dakota Howard, Dhwani Batra, Ben L. Rambo-Martin, Clinton R. Paden, Suxiang Tong, Duncan MacCannell |
| EPI_ISL_1073538, EPI_ISL_1210193, EPI_ISL_1210242, EPI_ISL_1210329 | Israel Central Virology laboratory | Israel National Consortium for SARS-CoV-2 sequencing | Neta Zuckerman, Efrat Dahan Bucris, Michal Mandelboim, Dana Bar-Ilan, Oran Erster, Tzvia Mann, Omer Murik, David A. Zeevi, Assaf Rokney, Joseph Jaffe, Eva Nachum, Maya Davidovich Cohen, Ephraim Fass, Gal Zizelski Valenci, Mor Rubinstein, Efrat Rorman, Israel Nissan, Efrat Glick-Saar, Omri Nayshool, Gideon Rechavi, Ella Mendelson, Orna Mor |
| EPI_ISL_1751614 | Max von Pettenkofer Institute, Virology, National Reference Center for Retroviruses, LMU Munich | Laboratory for Functional Genome Analysis; Dept. Genomics; Gene Center of the LMU Munich | Max Muenchhoff; Stefan Krebs; Alexander Graf; Oliver Keppler; Helmut Blum |
| EPI_ISL_1793836 | Hong Kong Department of Health | School of Public Health, The University of Hong Kong | Dominic N.C. Tsang, Daniel K.W. Chu, Haogao Gu, Tong Zhang, Leo L.M. Poon, Malik Peiris |
| EPI_ISL_1867110 | Department of Virus and Microbiological Special Diagnostics, Statens Serum Institut, Copenhagen, Denmark | Aalborg University | Danish Covid-19 Genome Consortium |
| EPI_ISL_566194 | Lighthouse Lab in Alderley Park | Wellcome Sanger Institute for the COVID-19 Genomics UK (COG-UK) consortium | Jacquelyn Wynn, Mairead Hyland, The Lighthouse Lab in Alderley Park and Alex Alderton, Roberto Amato, Sonia Goncalves, Ewan Harrison, David K. Jackson, Ian Johnston, Dominic Kwiatkowski, Cordelia Langford, John Sillitoe on behalf of the Wellcome Sanger Institute COVID-19 Surveillance Team |
| EPI_ISL_567085, EPI_ISL_567105, EPI_ISL_567258 | Lighthouse Lab in Glasgow | Wellcome Sanger Institute for the COVID-19 Genomics UK (COG-UK) consortium | Harper VanSteenhouse, Yumi Kasai, David Gray, Carol Clugston, Anna Dominiczak and Alex Alderton, Roberto Amato, Sonia Goncalves, Ewan Harrison, David K. Jackson, Ian Johnston, Dominic Kwiatkowski, Cordelia Langford, John Sillitoe on behalf of the Wellcome Sanger Institute COVID-19 Surveillance Team |
| EPI_ISL_567541 | Lighthouse Lab in Alderley Park | Wellcome Sanger Institute for the COVID-19 Genomics UK (COG-UK) consortium | Jacquelyn Wynn, Mairead Hyland, The Lighthouse Lab in Alderley Park and Alex Alderton, Roberto Amato, Sonia Goncalves, Ewan Harrison, David K. Jackson, Ian Johnston, Dominic Kwiatkowski, Cordelia Langford, John Sillitoe on behalf of the Wellcome Sanger Institute COVID-19 Surveillance Team |
| EPI_ISL_572658 | Quadram Institute Bioscience | COVID-19 Genomics UK (COG-UK) Consortium | Dave J. Baker, Gemma L. Kay, Alp Aydin, Thanh Le-Viet, Steven Rudder, Ana P. Tedim, Anastasia Kolyva, Maria Diaz, Leonardo de Oliveira Martins, Nabil-Fareed Alikhan, Lizzie Meadows, Rachael Stanley, Ngozi Elumogo, Muhammed Yasir, Nicholas M. Thomson, Alexander J Trotter, Rachel Gilroy, Samuel Bloomfield, Claire Stuart, Andrew Bell, Reenesh Prakash, Samir Dervisevic, Alison E. Mather, John Wain, Mark Webber, Andrew J. Page, Justin O'Grady |
| EPI_ISL_575332 | Israel Central Virology laboratory | Israel Central Virology laboratory | Neta Zuckerman, Efrat Dahan Bucris, Oran Erster, Ella Mendelson, Michal Mandelboim |
| EPI_ISL_575461, EPI_ISL_575760 | Lighthouse Lab in Alderley Park | Wellcome Sanger Institute for the COVID-19 Genomics UK (COG-UK) consortium | Jacquelyn Wynn, Mairead Hyland, The Lighthouse Lab in Alderley Park and Alex Alderton, Roberto Amato, Sonia Goncalves, Ewan Harrison, David K. Jackson, Ian Johnston, Dominic Kwiatkowski, Cordelia Langford, John Sillitoe on behalf of the Wellcome Sanger Institute COVID-19 Surveillance Team |
| EPI_ISL_575983 | Lighthouse Lab in Alderley Park | Wellcome Sanger Institute for the COVID-19 Genomics UK (COG-UK) consortium | Jacquelyn Wynn, Mairead Hyland, The Lighthouse Lab in Alderley Park and Alex Alderton, Roberto Amato, Sonia Goncalves, Ewan Harrison, David K. Jackson, Ian Johnston, Dominic Kwiatkowski, Cordelia Langford, John Sillitoe on behalf of the Wellcome Sanger Institute COVID-19 Surveillance Team (http://www.sanger.ac.uk/covid-team) |
| EPI_ISL_579857 | Lighthouse Lab in Glasgow | Wellcome Sanger Institute for the COVID-19 Genomics UK (COG-UK) consortium | Harper VanSteenhouse, Yumi Kasai, David Gray, Carol Clugston, Anna Dominiczak and Alex Alderton, Roberto Amato, Sonia Goncalves, Ewan Harrison, David K. Jackson, Ian Johnston, Dominic Kwiatkowski, Cordelia Langford, John Sillitoe on behalf of the Wellcome Sanger Institute COVID-19 Surveillance Team |
| EPI_ISL_580052, EPI_ISL_580230 | Lighthouse Lab in Alderley Park | Wellcome Sanger Institute for the COVID-19 Genomics UK (COG-UK) consortium | Jacquelyn Wynn, Mairead Hyland, The Lighthouse Lab in Alderley Park and Alex Alderton, Roberto Amato, Sonia Goncalves, Ewan Harrison, David K. Jackson, Ian Johnston, Dominic Kwiatkowski, Cordelia Langford, John Sillitoe on behalf of the Wellcome Sanger Institute COVID-19 Surveillance Team |
| EPI_ISL_580453 | Lighthouse Lab in Cambridge | Wellcome Sanger Institute for the COVID-19 Genomics UK (COG-UK) consortium | Rob Howes, The Lighthouse Lab in Cambridge and Alex Alderton, Roberto Amato, Sonia Goncalves, Ewan Harrison, David K. Jackson, Ian Johnston, Dominic Kwiatkowski, Cordelia Langford, John Sillitoe on behalf of the Wellcome Sanger Institute COVID-19 Surveillance Team |
| EPI_ISL_580628 | Lighthouse Lab in Alderley Park | Wellcome Sanger Institute for the COVID-19 Genomics UK (COG-UK) consortium | Jacquelyn Wynn, Mairead Hyland, The Lighthouse Lab in Alderley Park and Alex Alderton, Roberto Amato, Sonia Goncalves, Ewan Harrison, David K. Jackson, Ian Johnston, Dominic Kwiatkowski, Cordelia Langford, John Sillitoe on behalf of the Wellcome Sanger Institute COVID-19 Surveillance Team |
| EPI_ISL_584384 | Queens Medical Centre, Clinical Microbiology Department / DeepSeq Nottingham | COVID-19 Genomics UK (COG-UK) Consortium | Gemma Clark, Wendy Smith, Manjinder Khakh, Vicki M Fleming, Michelle M Lister, Hannah Howson-Wells, Jonathan Ball, Patrick McClure, Joseph Chappell, Theocharis Tsoleridis, Nadine Holmes, Matthew Carlisle, Christopher Moore, Fei Sang, Johnny Debebe, Victoria Wright, Matthew Loose |
| EPI_ISL_585479 | Virology Department, Royal Infirmary of Edinburgh, NHS Lothian / School of Biological Sciences, University of Edinburgh / Institute of Genetics and Molecular Medicine, University of Edinburgh | COVID-19 Genomics UK (COG-UK) Consortium | McHugh M, Dewar R, Rooke S, Gallagher M, Balcaza C, O'Toole Á, Scher E, Hill V, McCrone JT, Colquhoun R, Yu X, Jackson B, Rambaut A, Williams TC, Templeton K |
| EPI_ISL_586691, EPI_ISL_586953 | Lighthouse Lab in Alderley Park | Wellcome Sanger Institute for the COVID-19 Genomics UK (COG-UK) consortium | Jacquelyn Wynn, Mairead Hyland, The Lighthouse Lab in Alderley Park and Alex Alderton, Roberto Amato, Sonia Goncalves, Ewan Harrison, David K. Jackson, Ian Johnston, Dominic Kwiatkowski, Cordelia Langford, John Sillitoe on behalf of the Wellcome Sanger Institute COVID-19 Surveillance Team (http://www.sanger.ac.uk/covid-team) |
| EPI_ISL_587181, EPI_ISL_587252, EPI_ISL_587735 | Lighthouse Lab in Alderley Park | Wellcome Sanger Institute for the COVID-19 Genomics UK (COG-UK) consortium | Jacquelyn Wynn, Mairead Hyland, The Lighthouse Lab in Alderley Park and Alex Alderton, Roberto Amato, Sonia Goncalves, Ewan Harrison, David K. Jackson, Ian Johnston, Dominic Kwiatkowski, Cordelia Langford, John Sillitoe on behalf of the Wellcome Sanger Institute COVID-19 Surveillance Team |
| EPI_ISL_588330, EPI_ISL_588594, EPI_ISL_588980, EPI_ISL_589192 | Lighthouse Lab in Glasgow | Wellcome Sanger Institute for the COVID-19 Genomics UK (COG-UK) consortium | Harper VanSteenhouse, Yumi Kasai, David Gray, Carol Clugston, Anna Dominiczak and Alex Alderton, Roberto Amato, Sonia Goncalves, Ewan Harrison, David K. Jackson, Ian Johnston, Dominic Kwiatkowski, Cordelia Langford, John Sillitoe on behalf of the Wellcome Sanger Institute COVID-19 Surveillance Team |
| EPI_ISL_597229, EPI_ISL_597308 | Lighthouse Lab in Cambridge | Wellcome Sanger Institute for the COVID-19 Genomics UK (COG-UK) consortium | Rob Howes, The Lighthouse Lab in Cambridge and Alex Alderton, Roberto Amato, Sonia Goncalves, Ewan Harrison, David K. Jackson, Ian Johnston, Dominic Kwiatkowski, Cordelia Langford, John Sillitoe on behalf of the Wellcome Sanger Institute COVID-19 Surveillance Team (http://www.sanger.ac.uk/covid-team) |
| EPI_ISL_597667, EPI_ISL_598060, EPI_ISL_598063, EPI_ISL_598082, EPI_ISL_598083, EPI_ISL_598109, EPI_ISL_598172, EPI_ISL_598228 | Lighthouse Lab in Alderley Park | Wellcome Sanger Institute for the COVID-19 Genomics UK (COG-UK) consortium | Jacquelyn Wynn, Mairead Hyland, The Lighthouse Lab in Alderley Park and Alex Alderton, Roberto Amato, Sonia Goncalves, Ewan Harrison, David K. Jackson, Ian Johnston, Dominic Kwiatkowski, Cordelia Langford, John Sillitoe on behalf of the Wellcome Sanger Institute COVID-19 Surveillance Team (http://www.sanger.ac.uk/covid-team) |
| EPI_ISL_599524, EPI_ISL_599565, EPI_ISL_599612, EPI_ISL_600813 | Lighthouse Lab in Glasgow | Wellcome Sanger Institute for the COVID-19 Genomics UK (COG-UK) consortium | Harper VanSteenhouse, Yumi Kasai, David Gray, Carol Clugston, Anna Dominiczak and Alex Alderton, Roberto Amato, Sonia Goncalves, Ewan Harrison, David K. Jackson, Ian Johnston, Dominic Kwiatkowski, Cordelia Langford, John Sillitoe on behalf of the Wellcome Sanger Institute COVID-19 Surveillance Team (http://www.sanger.ac.uk/covid-team) |
| EPI_ISL_605954, EPI_ISL_606227 | Lighthouse Lab in Alderley Park | Wellcome Sanger Institute for the COVID-19 Genomics UK (COG-UK) consortium | Jacquelyn Wynn, Mairead Hyland, The Lighthouse Lab in Alderley Park and Alex Alderton, Roberto Amato, Sonia Goncalves, Ewan Harrison, David K. Jackson, Ian Johnston, Dominic Kwiatkowski, Cordelia Langford, John Sillitoe on behalf of the Wellcome Sanger Institute COVID-19 Surveillance Team |
| EPI_ISL_606472 | Lighthouse Lab in Milton Keynes | Wellcome Sanger Institute for the COVID-19 Genomics UK | The Lighthouse Lab in Milton Keynes and Alex Alderton, Roberto Amato, Sonia Goncalves, Ewan Harrison, David K. Jackson, Ian Johnston, Dominic |

| | | | |
|---|---|---|---|
| | | (COG-UK) consortium | Kwiatkowski, Cordelia Langford, John Sillitoe on behalf of the Wellcome Sanger Institute COVID-19 Surveillance Team |
| EPI_ISL_607059, EPI_ISL_607785, EPI_ISL_607998, EPI_ISL_608659, EPI_ISL_608663 | Lighthouse Lab in Alderley Park | Wellcome Sanger Institute for the COVID-19 Genomics UK (COG-UK) consortium | Jacquelyn Wynn, Mairead Hyland, The Lighthouse Lab in Alderley Park and Alex Alderton, Roberto Amato, Sonia Goncalves, Ewan Harrison, David K. Jackson, Ian Johnston, Dominic Kwiatkowski, Cordelia Langford, John Sillitoe on behalf of the Wellcome Sanger Institute COVID-19 Surveillance Team |
| EPI_ISL_608809 | Lighthouse Lab in Milton Keynes | Wellcome Sanger Institute for the COVID-19 Genomics UK (COG-UK) consortium | The Lighthouse Lab in Milton Keynes and Alex Alderton, Roberto Amato, Sonia Goncalves, Ewan Harrison, David K. Jackson, Ian Johnston, Dominic Kwiatkowski, Cordelia Langford, John Sillitoe on behalf of the Wellcome Sanger Institute COVID-19 Surveillance Team |
| EPI_ISL_609312, EPI_ISL_609362, EPI_ISL_609757 | Lighthouse Lab in Cambridge | Wellcome Sanger Institute for the COVID-19 Genomics UK (COG-UK) consortium | Rob Howes, The Lighthouse Lab in Cambridge and Alex Alderton, Roberto Amato, Sonia Goncalves, Ewan Harrison, David K. Jackson, Ian Johnston, Dominic Kwiatkowski, Cordelia Langford, John Sillitoe on behalf of the Wellcome Sanger Institute COVID-19 Surveillance Team |
| EPI_ISL_610342, EPI_ISL_610484 | Lighthouse Lab in Alderley Park | Wellcome Sanger Institute for the COVID-19 Genomics UK (COG-UK) consortium | Jacquelyn Wynn, Mairead Hyland, The Lighthouse Lab in Alderley Park and Alex Alderton, Roberto Amato, Sonia Goncalves, Ewan Harrison, David K. Jackson, Ian Johnston, Dominic Kwiatkowski, Cordelia Langford, John Sillitoe on behalf of the Wellcome Sanger Institute COVID-19 Surveillance Team |
| EPI_ISL_610659 | Lighthouse Lab in Cambridge | Wellcome Sanger Institute for the COVID-19 Genomics UK (COG-UK) consortium | Rob Howes, The Lighthouse Lab in Cambridge and Alex Alderton, Roberto Amato, Sonia Goncalves, Ewan Harrison, David K. Jackson, Ian Johnston, Dominic Kwiatkowski, Cordelia Langford, John Sillitoe on behalf of the Wellcome Sanger Institute COVID-19 Surveillance Team |
| EPI_ISL_611280 | Lighthouse Lab in Alderley Park | Wellcome Sanger Institute for the COVID-19 Genomics UK (COG-UK) consortium | Jacquelyn Wynn, Mairead Hyland, The Lighthouse Lab in Alderley Park and Alex Alderton, Roberto Amato, Sonia Goncalves, Ewan Harrison, David K. Jackson, Ian Johnston, Dominic Kwiatkowski, Cordelia Langford, John Sillitoe on behalf of the Wellcome Sanger Institute COVID-19 Surveillance Team (http://www.sanger.ac.uk/covid-team) |
| EPI_ISL_611479 | Lighthouse Lab in Glasgow | Wellcome Sanger Institute for the COVID-19 Genomics UK (COG-UK) consortium | Harper VanSteenhouse, Yumi Kasai, David Gray, Carol Clugston, Anna Dominiczak and Alex Alderton, Roberto Amato, Sonia Goncalves, Ewan Harrison, David K. Jackson, Ian Johnston, Dominic Kwiatkowski, Cordelia Langford, John Sillitoe on behalf of the Wellcome Sanger Institute COVID-19 Surveillance Team (http://www.sanger.ac.uk/covid-team) |
| EPI_ISL_623369, EPI_ISL_623534, EPI_ISL_623638 | Lighthouse Lab in Milton Keynes | Wellcome Sanger Institute for the COVID-19 Genomics UK (COG-UK) consortium | The Lighthouse Lab in Milton Keynes and Alex Alderton, Roberto Amato, Sonia Goncalves, Ewan Harrison, David K. Jackson, Ian Johnston, Dominic Kwiatkowski, Cordelia Langford, John Sillitoe on behalf of the Wellcome Sanger Institute COVID-19 Surveillance Team (http://www.sanger.ac.uk/covid-team) |
| EPI_ISL_623940, EPI_ISL_624411, EPI_ISL_624578, EPI_ISL_624780, EPI_ISL_624953, EPI_ISL_625027, EPI_ISL_625161, EPI_ISL_625238 | Lighthouse Lab in Alderley Park | Wellcome Sanger Institute for the COVID-19 Genomics UK (COG-UK) consortium | Jacquelyn Wynn, Mairead Hyland, The Lighthouse Lab in Alderley Park and Alex Alderton, Roberto Amato, Sonia Goncalves, Ewan Harrison, David K. Jackson, Ian Johnston, Dominic Kwiatkowski, Cordelia Langford, John Sillitoe on behalf of the Wellcome Sanger Institute COVID-19 Surveillance Team (http://www.sanger.ac.uk/covid-team) |
| EPI_ISL_625371 | Lighthouse Lab in Glasgow | Wellcome Sanger Institute for the COVID-19 Genomics UK (COG-UK) consortium | Harper VanSteenhouse, Yumi Kasai, David Gray, Carol Clugston, Anna Dominiczak and Alex Alderton, Roberto Amato, Sonia Goncalves, Ewan Harrison, David K. Jackson, Ian Johnston, Dominic Kwiatkowski, Cordelia Langford, John Sillitoe on behalf of the Wellcome Sanger Institute COVID-19 Surveillance Team (http://www.sanger.ac.uk/covid-team) |
| EPI_ISL_630022 | Lighthouse Lab in Milton Keynes | Wellcome Sanger Institute for the COVID-19 Genomics UK (COG-UK) consortium | The Lighthouse Lab in Milton Keynes and Alex Alderton, Roberto Amato, Sonia Goncalves, Ewan Harrison, David K. Jackson, Ian Johnston, Dominic Kwiatkowski, Cordelia Langford, John Sillitoe on behalf of the Wellcome Sanger Institute COVID-19 Surveillance Team |
| EPI_ISL_630354, EPI_ISL_630690, EPI_ISL_630889 | Lighthouse Lab in Alderley Park | Wellcome Sanger Institute for the COVID-19 Genomics UK (COG-UK) consortium | Jacquelyn Wynn, Mairead Hyland, The Lighthouse Lab in Alderley Park and Alex Alderton, Roberto Amato, Sonia Goncalves, Ewan Harrison, David K. Jackson, Ian Johnston, Dominic Kwiatkowski, Cordelia Langford, John Sillitoe on behalf of the Wellcome Sanger Institute COVID-19 Surveillance Team |
| EPI_ISL_633492 | Lighthouse Lab in Glasgow | Wellcome Sanger Institute for the COVID-19 Genomics UK (COG-UK) Consortium | Harper VanSteenhouse, Yumi Kasai, David Gray, Carol Clugston, Anna Dominiczak and Alex Alderton, Roberto Amato, Sonia Goncalves, Ewan Harrison, David K. Jackson, Ian Johnston, Dominic Kwiatkowski, Cordelia Langford, John Sillitoe on behalf of the Wellcome Sanger Institute COVID-19 Surveillance Team |
| EPI_ISL_634257 | Lighthouse Lab in Milton Keynes | Wellcome Sanger Institute for the COVID-19 Genomics UK (COG-UK) consortium | The Lighthouse Lab in Milton Keynes and Alex Alderton, Roberto Amato, Sonia Goncalves, Ewan Harrison, David K. Jackson, Ian Johnston, Dominic Kwiatkowski, Cordelia Langford, John Sillitoe on behalf of the Wellcome Sanger Institute COVID-19 Surveillance Team |
| EPI_ISL_637858, EPI_ISL_638843, EPI_ISL_638849 | Oxford Viromics, NDM, University of Oxford; Oxford University Hospitals; Basingstoke and North Hampshire Hospital | COVID-19 Genomics UK (COG-UK) Consortium | Tanya Golubchik, David Bonsall, George Macintyre, Amy Trebes, Mariateresa de Cesare, Catrin Moore, Alex Mobbs, Anita Justice, Robert Shaw, Monique Andersson, Timothy Peto, Emma Wise, Nathan Moore, Jessica Lynch, Nick Cortes, Matilde Mori, Stephen Kidd, David Buck, John Todd, Christophe Fraser |
| EPI_ISL_642252, EPI_ISL_642319, EPI_ISL_642350, EPI_ISL_642593, EPI_ISL_642628, EPI_ISL_643214, EPI_ISL_643268 | Lighthouse Lab in Glasgow | Wellcome Sanger Institute for the COVID-19 Genomics UK (COG-UK) Consortium | Harper VanSteenhouse, Yumi Kasai, David Gray, Carol Clugston, Anna Dominiczak and Alex Alderton, Roberto Amato, Sonia Goncalves, Ewan Harrison, David K. Jackson, Ian Johnston, Dominic Kwiatkowski, Cordelia Langford, John Sillitoe on behalf of the Wellcome Sanger Institute COVID-19 Surveillance Team |
| EPI_ISL_645539, EPI_ISL_645606, EPI_ISL_645722, EPI_ISL_646015, EPI_ISL_646166, EPI_ISL_646180, EPI_ISL_646327, EPI_ISL_646428 | Lighthouse Lab in Alderley Park | Wellcome Sanger Institute for the COVID-19 Genomics UK (COG-UK) Consortium | Jacquelyn Wynn, Mairead Hyland, The Lighthouse Lab in Alderley Park and Alex Alderton, Roberto Amato, Sonia Goncalves, Ewan Harrison, David K. Jackson, Ian Johnston, Dominic Kwiatkowski, Cordelia Langford, John Sillitoe on behalf of the Wellcome Sanger Institute COVID-19 Surveillance Team |
| EPI_ISL_646450 | Lighthouse Lab in Glasgow | Wellcome Sanger Institute for the COVID-19 Genomics UK (COG-UK) Consortium | Harper VanSteenhouse, Yumi Kasai, David Gray, Carol Clugston, Anna Dominiczak and Alex Alderton, Roberto Amato, Sonia Goncalves, Ewan Harrison, David K. Jackson, Ian Johnston, Dominic Kwiatkowski, Cordelia Langford, John Sillitoe on behalf of the Wellcome Sanger Institute COVID-19 Surveillance Team |
| EPI_ISL_646538, EPI_ISL_646638, EPI_ISL_646716, EPI_ISL_646755, EPI_ISL_647054, EPI_ISL_647071 | Lighthouse Lab in Alderley Park | Wellcome Sanger Institute for the COVID-19 Genomics UK (COG-UK) Consortium | Jacquelyn Wynn, Mairead Hyland, The Lighthouse Lab in Alderley Park and Alex Alderton, Roberto Amato, Sonia Goncalves, Ewan Harrison, David K. Jackson, Ian Johnston, Dominic Kwiatkowski, Cordelia Langford, John Sillitoe on behalf of the Wellcome Sanger Institute COVID-19 Surveillance Team |
| EPI_ISL_647279 | Lighthouse Lab in Cambridge | Wellcome Sanger Institute for the COVID-19 Genomics UK (COG-UK) Consortium | Rob Howes, The Lighthouse Lab in Cambridge and Alex Alderton, Roberto Amato, Sonia Goncalves, Ewan Harrison, David K. Jackson, Ian Johnston, Dominic Kwiatkowski, Cordelia Langford, John Sillitoe on behalf of the Wellcome Sanger Institute COVID-19 Surveillance Team |
| EPI_ISL_647511, EPI_ISL_647541, EPI_ISL_647776 | Lighthouse Lab in Alderley Park | Wellcome Sanger Institute for the COVID-19 Genomics UK (COG-UK) Consortium | Jacquelyn Wynn, Mairead Hyland, The Lighthouse Lab in Alderley Park and Alex Alderton, Roberto Amato, Sonia Goncalves, Ewan Harrison, David K. Jackson, Ian Johnston, Dominic Kwiatkowski, Cordelia Langford, John Sillitoe on behalf of the Wellcome Sanger Institute COVID-19 Surveillance Team |
| EPI_ISL_650816 | West of Scotland Specialist Virology Centre, NHSGGC / MRC-University of Glasgow Centre for Virus Research | COVID-19 Genomics UK (COG-UK) Consortium | Ana da Silva Filipe, Natasha Johnson, Kathy Smollett, Daniel Mair, Stephen Carmichael, Alice Broos, Lily Tong, Jenna Nichols, Kyriaki Nomikou; Sarah McDonald; Richard Orton, Joseph Hughes, Sreenu Vattipally, David L Robertson; Alasdair MacLean, Rory Gunson; Sharif Shaaban, Matthew Holden; Rachel Blacow, Guy Mollett, Kathy Li, James Shepherd, Antonia Ho, Emma Thomson |
| EPI_ISL_651372, EPI_ISL_652691, EPI_ISL_652695 | University of Exeter | COVID-19 Genomics UK (COG-UK) Consortium | Ben Temperton,Aaron Jeffries,Michelle Michelsen,Joanna Warwick-Dugdale,Audrey Farbos,Robyn Manley,Stephen Michell,Jane Masoli |
| EPI_ISL_652782 | Oxford Viromics, NDM, University of Oxford; Oxford University Hospitals; Basingstoke and North Hampshire Hospital | COVID-19 Genomics UK (COG-UK) Consortium | Tanya Golubchik, David Bonsall, George Macintyre, Amy Trebes, Mariateresa de Cesare, Catrin Moore, Alex Mobbs, Anita Justice, Robert Shaw, Monique Andersson, Timothy Peto, Emma Wise, Nathan Moore, Jessica Lynch, Nick Cortes, Matilde Mori, Stephen Kidd, David Buck, John Todd, Christophe Fraser |
| EPI_ISL_655830 | Lighthouse Lab in Alderley Park | Wellcome Sanger Institute for the COVID-19 Genomics UK (COG-UK) Consortium | Jacquelyn Wynn, Mairead Hyland, The Lighthouse Lab in Alderley Park and Alex Alderton, Roberto Amato, Sonia Goncalves, Ewan Harrison, David K. Jackson, Ian Johnston, Dominic Kwiatkowski, Cordelia Langford, John Sillitoe on behalf of the Wellcome Sanger Institute COVID-19 Surveillance Team |
| EPI_ISL_655918 | Lighthouse Lab in Glasgow | Wellcome Sanger Institute for the COVID-19 Genomics UK (COG-UK) Consortium | Harper VanSteenhouse, Yumi Kasai, David Gray, Carol Clugston, Anna Dominiczak and Alex Alderton, Roberto Amato, Sonia Goncalves, Ewan Harrison, David K. Jackson, Ian Johnston, Dominic Kwiatkowski, Cordelia Langford, John Sillitoe on behalf of the Wellcome Sanger Institute COVID-19 Surveillance Team |
| EPI_ISL_656146, EPI_ISL_656210, EPI_ISL_656328 | Lighthouse Lab in Alderley Park | Wellcome Sanger Institute for the COVID-19 Genomics UK (COG-UK) Consortium | Jacquelyn Wynn, Mairead Hyland, The Lighthouse Lab in Alderley Park and Alex Alderton, Roberto Amato, Sonia Goncalves, Ewan Harrison, David K. Jackson, Ian Johnston, Dominic Kwiatkowski, Cordelia Langford, John Sillitoe on behalf of the Wellcome Sanger Institute COVID-19 Surveillance Team |
| EPI_ISL_658583, EPI_ISL_658723 | Lighthouse Lab in Glasgow | Wellcome Sanger Institute for the COVID-19 Genomics UK (COG-UK) Consortium | Harper VanSteenhouse, Yumi Kasai, David Gray, Carol Clugston, Anna Dominiczak and Alex Alderton, Roberto Amato, Sonia Goncalves, Ewan Harrison, David K. Jackson, Ian Johnston, Dominic Kwiatkowski, Cordelia Langford, John Sillitoe on behalf of the Wellcome Sanger Institute COVID-19 Surveillance |

| | | | |
|---|---|---|---|
| | | | Team |
| EPI_ISL_658789 | Lighthouse Lab in Milton Keynes | Wellcome Sanger Institute for the COVID-19 Genomics UK (COG-UK) Consortium | The Lighthouse Lab in Milton Keynes and Alex Alderton, Roberto Amato, Sonia Goncalves, Ewan Harrison, David K. Jackson, Ian Johnston, Dominic Kwiatkowski, Cordelia Langford, John Sillitoe on behalf of the Wellcome Sanger Institute COVID-19 Surveillance Team |
| EPI_ISL_660044 | Lighthouse Lab in Glasgow | Wellcome Sanger Institute for the COVID-19 Genomics UK (COG-UK) Consortium | Harper VanSteenhouse, Yumi Kasai, David Gray, Carol Clugston, Anna Dominiczak and Alex Alderton, Roberto Amato, Sonia Goncalves, Ewan Harrison, David K. Jackson, Ian Johnston, Dominic Kwiatkowski, Cordelia Langford, John Sillitoe on behalf of the Wellcome Sanger Institute COVID-19 Surveillance Team (http://www.sanger.ac.uk/covid-team) |
| EPI_ISL_660764 | Respiratory Virus Unit, Microbiology Services Colindale, Public Health England | COVID-19 Genomics UK (COG-UK) Consortium | PHE Covid Sequencing Team |
| EPI_ISL_661400, EPI_ISL_662007 | Lighthouse Lab in Glasgow | Wellcome Sanger Institute for the COVID-19 Genomics UK (COG-UK) Consortium | Harper VanSteenhouse, Yumi Kasai, David Gray, Carol Clugston, Anna Dominiczak and Alex Alderton, Roberto Amato, Sonia Goncalves, Ewan Harrison, David K. Jackson, Ian Johnston, Dominic Kwiatkowski, Cordelia Langford, John Sillitoe on behalf of the Wellcome Sanger Institute COVID-19 Surveillance Team |
| EPI_ISL_662029, EPI_ISL_662133 | Lighthouse Lab in Alderley Park | Wellcome Sanger Institute for the COVID-19 Genomics UK (COG-UK) Consortium | Jacquelyn Wynn, Mairead Hyland, The Lighthouse Lab in Alderley Park and Alex Alderton, Roberto Amato, Sonia Goncalves, Ewan Harrison, David K. Jackson, Ian Johnston, Dominic Kwiatkowski, Cordelia Langford, John Sillitoe on behalf of the Wellcome Sanger Institute COVID-19 Surveillance Team |
| EPI_ISL_662188, EPI_ISL_662195, EPI_ISL_662729, EPI_ISL_662992 | Lighthouse Lab in Glasgow | Wellcome Sanger Institute for the COVID-19 Genomics UK (COG-UK) Consortium | Harper VanSteenhouse, Yumi Kasai, David Gray, Carol Clugston, Anna Dominiczak and Alex Alderton, Roberto Amato, Sonia Goncalves, Ewan Harrison, David K. Jackson, Ian Johnston, Dominic Kwiatkowski, Cordelia Langford, John Sillitoe on behalf of the Wellcome Sanger Institute COVID-19 Surveillance Team |
| EPI_ISL_664430 | Liverpool Clinical Laboratories | COVID-19 Genomics UK (COG-UK) Consortium | Sam Haldenby, Anita Lucaci, Steve Paterson, Julian Hiscox, Alistair Darby, M Almsaud, A Alrezaihi, Muhannad Alruwaili, Stuart D Armstrong, Jones Benjamin, Eleanor G Bentley, Anu Chawla, Jordan J Clark, Angela Cowell, Richard Eccles, Isabel García-Dorival, Matthew Gemmell, Alessandro Gerada, PKF Gilmore, Richard Gregory, Ximeng Han, Catherine Hartley, Margaret Hughes, Miren Iturriza-Gomara, James Johnson, L Luu, Jenifer Manson, Charlotte Nelson, Elaine O'Toole, Cassie Olateju, Rebekah Penrice-Randal , Lucille Rainbow, N.P Randle, Trevor Ian Robinson, Parul Sharma, Ghada T Shawli, James P Stewart, Neil Swainston, Ecaterina Vamos, Joanne Watts, Mark Whitehead |
| EPI_ISL_666729, EPI_ISL_666731, EPI_ISL_666732, EPI_ISL_666733, EPI_ISL_666734, EPI_ISL_666735 | Respiratory Virus Unit, Microbiology Services Colindale, Public Health England | COVID-19 Genomics UK (COG-UK) Consortium | PHE Covid Sequencing Team |
| EPI_ISL_667868, EPI_ISL_667940 | Lighthouse Lab in Glasgow | Wellcome Sanger Institute for the COVID-19 Genomics UK (COG-UK) Consortium | Harper VanSteenhouse, Yumi Kasai, David Gray, Carol Clugston, Anna Dominiczak and Alex Alderton, Roberto Amato, Sonia Goncalves, Ewan Harrison, David K. Jackson, Ian Johnston, Dominic Kwiatkowski, Cordelia Langford, John Sillitoe on behalf of the Wellcome Sanger Institute COVID-19 Surveillance Team |
| EPI_ISL_672996, EPI_ISL_673530 | Lighthouse Lab in Cambridge | Wellcome Sanger Institute for the COVID-19 Genomics UK (COG-UK) Consortium | Rob Howes, The Lighthouse Lab in Cambridge and Alex Alderton, Roberto Amato, Sonia Goncalves, Ewan Harrison, David K. Jackson, Ian Johnston, Dominic Kwiatkowski, Cordelia Langford, John Sillitoe on behalf of the Wellcome Sanger Institute COVID-19 Surveillance Team |
| EPI_ISL_674065 | Lighthouse Lab in Milton Keynes | Wellcome Sanger Institute for the COVID-19 Genomics UK (COG-UK) Consortium | The Lighthouse Lab in Milton Keynes and Alex Alderton, Roberto Amato, Sonia Goncalves, Ewan Harrison, David K. Jackson, Ian Johnston, Dominic Kwiatkowski, Cordelia Langford, John Sillitoe on behalf of the Wellcome Sanger Institute COVID-19 Surveillance Team |
| EPI_ISL_675706 | Lighthouse Lab in Alderley Park | Wellcome Sanger Institute for the COVID-19 Genomics UK (COG-UK) Consortium | Jacquelyn Wynn, Mairead Hyland, The Lighthouse Lab in Alderley Park and Alex Alderton, Roberto Amato, Sonia Goncalves, Ewan Harrison, David K. Jackson, Ian Johnston, Dominic Kwiatkowski, Cordelia Langford, John Sillitoe on behalf of the Wellcome Sanger Institute COVID-19 Surveillance Team |
| EPI_ISL_676431 | Lighthouse Lab in Glasgow | Wellcome Sanger Institute for the COVID-19 Genomics UK (COG-UK) Consortium | Harper VanSteenhouse, Yumi Kasai, David Gray, Carol Clugston, Anna Dominiczak and Alex Alderton, Roberto Amato, Sonia Goncalves, Ewan Harrison, David K. Jackson, Ian Johnston, Dominic Kwiatkowski, Cordelia Langford, John Sillitoe on behalf of the Wellcome Sanger Institute COVID-19 Surveillance Team |
| EPI_ISL_678662 | Respiratory Virus Unit, Microbiology Services Colindale, Public Health England | COVID-19 Genomics UK (COG-UK) Consortium | PHE Covid Sequencing Team |
| EPI_ISL_678883 | Department of Pathology, University of Cambridge | COVID-19 Genomics UK (COG-UK) Consortium | Aminu S. Jahun, Yasmin Chaudhry, Grant Hall, Iliana Georgana, Myra Hosmillo, Martin D. Curran, Malte Pinckert, Surendra Parmar, Ian Goodfellow |
| EPI_ISL_678926 | Queens Medical Centre, Clinical Microbiology Department / DeepSeq Nottingham | COVID-19 Genomics UK (COG-UK) Consortium | Gemma Clark, Wendy Smith, Manjinder Khakh, Vicki M Fleming, Michelle M Lister, Hannah Howson-Wells, Jonathan Ball, Patrick McClure, Joseph Chappell, Theocharis Tsoleridis, Nadine Holmes, Matthew Carlisle, Christopher Moore, Fei Sang, Johnny Debebe, Victoria Wright, Matthew Loose |
| EPI_ISL_679727, EPI_ISL_679728 | Oxford Viromics, NDM, University of Oxford; Oxford University Hospitals; Basingstoke and North Hampshire Hospital | COVID-19 Genomics UK (COG-UK) Consortium | Tanya Golubchik, David Bonsall, George Macintyre, Amy Trebes, Mariateresa de Cesare, Catrin Moore, Alex Mobbs, Anita Justice, Robert Shaw, Monique Andersson, Timothy Peto, Emma Wise, Nathan Moore, Jessica Lynch, Nick Cortes, Matilde Mori, Stephen Kidd, David Buck, John Todd, Christophe Fraser |
| EPI_ISL_701905, EPI_ISL_701996, EPI_ISL_702604, EPI_ISL_702813 | Lighthouse Lab in Alderley Park | Wellcome Sanger Institute for the COVID-19 Genomics UK (COG-UK) Consortium | Jacquelyn Wynn, Mairead Hyland, The Lighthouse Lab in Alderley Park and Alex Alderton, Roberto Amato, Sonia Goncalves, Ewan Harrison, David K. Jackson, Ian Johnston, Dominic Kwiatkowski, Cordelia Langford, John Sillitoe on behalf of the Wellcome Sanger Institute COVID-19 Surveillance Team |
| EPI_ISL_706211 | Oxford Viromics, NDM, University of Oxford; Oxford University Hospitals; Basingstoke and North Hampshire Hospital | COVID-19 Genomics UK (COG-UK) Consortium | Tanya Golubchik, David Bonsall, George Macintyre, Amy Trebes, Mariateresa de Cesare, Catrin Moore, Alex Mobbs, Anita Justice, Robert Shaw, Monique Andersson, Timothy Peto, Emma Wise, Nathan Moore, Jessica Lynch, Nick Cortes, Matilde Mori, Stephen Kidd, David Buck, John Todd, Christophe Fraser |
| EPI_ISL_709868 | Lighthouse Lab in Milton Keynes | Wellcome Sanger Institute for the COVID-19 Genomics UK (COG-UK) Consortium | The Lighthouse Lab in Milton Keynes and Alex Alderton, Roberto Amato, Sonia Goncalves, Ewan Harrison, David K. Jackson, Ian Johnston, Dominic Kwiatkowski, Cordelia Langford, John Sillitoe on behalf of the Wellcome Sanger Institute COVID-19 Surveillance Team |
| EPI_ISL_719666, EPI_ISL_719752, EPI_ISL_719782 | Lighthouse Lab in Glasgow | Wellcome Sanger Institute for the COVID-19 Genomics UK (COG-UK) Consortium | Harper VanSteenhouse, Yumi Kasai, David Gray, Carol Clugston, Anna Dominiczak and Alex Alderton, Roberto Amato, Sonia Goncalves, Ewan Harrison, David K. Jackson, Ian Johnston, Dominic Kwiatkowski, Cordelia Langford, John Sillitoe on behalf of the Wellcome Sanger Institute COVID-19 Surveillance Team |
| EPI_ISL_720381, EPI_ISL_720409 | Lighthouse Lab in Milton Keynes | Wellcome Sanger Institute for the COVID-19 Genomics UK (COG-UK) Consortium | The Lighthouse Lab in Milton Keynes and Alex Alderton, Roberto Amato, Sonia Goncalves, Ewan Harrison, David K. Jackson, Ian Johnston, Dominic Kwiatkowski, Cordelia Langford, John Sillitoe on behalf of the Wellcome Sanger Institute COVID-19 Surveillance Team |
| EPI_ISL_721014 | Lighthouse Lab in Glasgow | Wellcome Sanger Institute for the COVID-19 Genomics UK (COG-UK) Consortium | Harper VanSteenhouse, Yumi Kasai, David Gray, Carol Clugston, Anna Dominiczak and Alex Alderton, Roberto Amato, Sonia Goncalves, Ewan Harrison, David K. Jackson, Ian Johnston, Dominic Kwiatkowski, Cordelia Langford, John Sillitoe on behalf of the Wellcome Sanger Institute COVID-19 Surveillance Team |
| EPI_ISL_727788, EPI_ISL_727796 | Centre for Enzyme Innovation, University of Portsmouth / Translational Research Laboratory, Portsmouth Hospitals NHS Trust | COVID-19 Genomics UK (COG-UK) Consortium | Angela Beckett,Yann Bourgeois,Garry Scarlett,Sharon Glaysher,Scott Elliott,Kelly Bicknell,Robert Impey,Allyson Lloyd,Sarah Wyllie,Ethan Butcher,Anoop Chauhan,Samuel Robson |
| EPI_ISL_730784 | Lighthouse Lab in Alderley Park | Wellcome Sanger Institute for the COVID-19 Genomics UK (COG-UK) Consortium | Jacquelyn Wynn, Mairead Hyland, The Lighthouse Lab in Alderley Park and Alex Alderton, Roberto Amato, Sonia Goncalves, Ewan Harrison, David K. Jackson, Ian Johnston, Dominic Kwiatkowski, Cordelia Langford, John Sillitoe on behalf of the Wellcome Sanger Institute COVID-19 Surveillance Team |
| EPI_ISL_733578 | Temporary Specimen Collection Centre | Hong Kong Department of Health | Alan K.L. Tsang, Peter C.W. Yip, Edman T.K. Lam, Rickjason C.W. Chan, Dominic N.C. Tsang |
| EPI_ISL_733795 | Lighthouse Lab in Alderley Park | Wellcome Sanger Institute for the COVID-19 Genomics UK (COG-UK) Consortium | Jacquelyn Wynn, Mairead Hyland, The Lighthouse Lab in Alderley Park and Alex Alderton, Roberto Amato, Sonia Goncalves, Ewan Harrison, David K. Jackson, Ian Johnston, Dominic Kwiatkowski, Cordelia Langford, John Sillitoe on behalf of the Wellcome Sanger Institute COVID-19 Surveillance Team |
| EPI_ISL_740923 | Oxford Viromics, NDM, University of Oxford; Oxford University Hospitals; Basingstoke and North Hampshire Hospital | COVID-19 Genomics UK (COG-UK) Consortium | Tanya Golubchik, David Bonsall, George Macintyre, Amy Trebes, Mariateresa de Cesare, Catrin Moore, Alex Mobbs, Anita Justice, Robert Shaw, Monique Andersson, Timothy Peto, Emma Wise, Nathan Moore, Jessica Lynch, Nick Cortes, Matilde Mori, Stephen Kidd, David Buck, John Todd, Christophe Fraser |
| EPI_ISL_760770 | Lighthouse Lab in Glasgow | Wellcome Sanger Institute for the COVID-19 Genomics UK (COG-UK) Consortium | Harper VanSteenhouse, Yumi Kasai, David Gray, Carol Clugston, Anna Dominiczak and Alex Alderton, Roberto Amato, Sonia Goncalves, Ewan Harrison, David K. Jackson, Ian Johnston, Dominic Kwiatkowski, Cordelia Langford, John Sillitoe on behalf of the Wellcome Sanger Institute COVID-19 Surveillance Team |
| EPI_ISL_760949 | Lighthouse Lab in Alderley Park | Wellcome Sanger Institute for the COVID-19 Genomics UK (COG-UK) Consortium | Jacquelyn Wynn, Mairead Hyland, The Lighthouse Lab in Alderley Park and Alex Alderton, Roberto Amato, Sonia Goncalves, Ewan Harrison, David K. Jackson, Ian Johnston, Dominic Kwiatkowski, Cordelia Langford, John Sillitoe on behalf of the Wellcome Sanger Institute COVID-19 Surveillance Team |

| | | | |
|---|---|---|---|
| EPI_ISL_769500 | Lighthouse Lab in Glasgow | Wellcome Sanger Institute for the COVID-19 Genomics UK (COG-UK) Consortium | Harper VanSteenhouse, Yumi Kasai, David Gray, Carol Clugston, Anna Dominiczak and Alex Alderton, Roberto Amato, Sonia Goncalves, Ewan Harrison, David K. Jackson, Ian Johnston, Dominic Kwiatkowski, Cordelia Langford, John Sillitoe on behalf of the Wellcome Sanger Institute COVID-19 Surveillance Team |
| EPI_ISL_776586, EPI_ISL_776635, EPI_ISL_776828 | Israel Central Virology laboratory | Israel Central Virology laboratory | Neta Zuckerman, Efrat Dahan Bucris, Oran Erster, Michal Mandelboim, Orna Mor, Ella Mendelson |
| EPI_ISL_782402 | Lighthouse Lab in Alderley Park | Wellcome Sanger Institute for the COVID-19 Genomics UK (COG-UK) Consortium | Jacquelyn Wynn, Mairead Hyland, The Lighthouse Lab in Alderley Park and Alex Alderton, Roberto Amato, Sonia Goncalves, Ewan Harrison, David K. Jackson, Ian Johnston, Dominic Kwiatkowski, Cordelia Langford, John Sillitoe on behalf of the Wellcome Sanger Institute COVID-19 Surveillance Team |
| EPI_ISL_795420 | Department of Virus and Microbiological Special Diagnostics, Statens Serum Institut, Copenhagen, Denmark | Albertsen Lab, Department of Chemistry and Bioscience, Aalborg University, Denmark | Danish Covid-19 Genome Consortium |
| EPI_ISL_804108, EPI_ISL_804168, EPI_ISL_814139, EPI_ISL_814153, EPI_ISL_814234 | Israel Central Virology laboratory | Israel National Consortium for SARS-CoV-2 sequencing | Neta Zuckerman, Efrat Dahan Bucris, Michal Mandelboim, Dana Bar-Ilan, Oran Erster, Tzvia Mann, Omer Murik, David A. Zeevi, Assaf Rokney, Joseph Jaffe, Eva Nachum, Maya Davidovich Cohen, Ephraim Fass, Gal Zizelski Valenci, Mor Rubinstein, Efrat Rorman, Israel Nissan, Efrat Glick-Saar, Omri Nayshool, Gideon Rechavi, Ella Mendelson, Orna Mor |
| EPI_ISL_817537, EPI_ISL_817540 | Department of Virus and Microbiological Special Diagnostics, Statens Serum Institut, Copenhagen, Denmark | Albertsen Lab, Department of Chemistry and Bioscience, Aalborg University, Denmark | Danish Covid-19 Genome Consortium |
| EPI_ISL_821288, EPI_ISL_834081, EPI_ISL_834145 | Lighthouse Lab in Alderley Park | Wellcome Sanger Institute for the COVID-19 Genomics UK (COG-UK) Consortium | Jacquelyn Wynn, Mairead Hyland, The Lighthouse Lab in Alderley Park and Alex Alderton, Roberto Amato, Sonia Goncalves, Ewan Harrison, David K. Jackson, Ian Johnston, Dominic Kwiatkowski, Cordelia Langford, John Sillitoe on behalf of the Wellcome Sanger Institute COVID-19 Surveillance Team |
| EPI_ISL_846908 | Department of Virus and Microbiological Special Diagnostics, Statens Serum Institut, Copenhagen, Denmark | Albertsen Lab, Department of Chemistry and Bioscience, Aalborg University, Denmark | Danish Covid-19 Genome Consortium |
| EPI_ISL_852593 | Max von Pettenkofer Institute, Virology, National Reference Center for Retroviruses, LMU München | Laboratory for Functional Genome Analysis, Dept. Genomics, Gene Center of the LMU Munich | Max Muenchhoff, Stefan Krebs, Alexander Graf, Oliver Keppler, Helmut Blum |
| EPI_ISL_863233, EPI_ISL_864126 | Lighthouse Lab in Alderley Park | Wellcome Sanger Institute for the COVID-19 Genomics UK (COG-UK) Consortium | Jacquelyn Wynn, Mairead Hyland, The Lighthouse Lab in Alderley Park and Alex Alderton, Roberto Amato, Sonia Goncalves, Ewan Harrison, David K. |
| EPI_ISL_865442 | Liverpool Clinical Laboratories | COVID-19 Genomics UK (COG-UK) Consortium | Sam Haldenby, Anita Lucaci, Steve Paterson, Julian Hiscox, Alistair Darby, M Almsaud, A Alrezaihi, Muhannad Alruwaili, Stuart D Armstrong, Jones Benjamin, Eleanor G Bentley, Anu Chawla, Jordan J Clark, Angela Cowell, Richard Eccles, Isabel García-Dorival, Matthew Gemmell, Alessandro Gerada, PKF Gilmore, Richard Gregory, Ximeng Han, Catherine Hartley, Margaret Hughes, Miren Iturriza-Gomara, James Johnson, L Luu, Jenifer Manson, Charlotte Nelson, Elaine O'Toole, Cassie Olateju, Rebekah Penrice-Randal , Lucille Rainbow, N.P Randle, Trevor Ian Robinson, Parul Sharma, Ghada T Shawli, James P Stewart, Neil Swainston, Ecaterina Vamos, Joanne Watts, Mark Whitehead |
| EPI_ISL_869688 | Department of Virus and Microbiological Special Diagnostics, Statens Serum Institut, Copenhagen, Denmark | Aalborg University | Danish Covid-19 Genome Consortium |
| EPI_ISL_889028, EPI_ISL_889185 | Israel Central Virology laboratory | Israel National Consortium for SARS-CoV-2 sequencing | Neta Zuckerman, Efrat Dahan Bucris, Michal Mandelboim, Dana Bar-Ilan, Oran Erster, Tzvia Mann, Omer Murik, David A. Zeevi, Assaf Rokney, Joseph Jaffe, Eva Nachum, Maya Davidovich Cohen, Ephraim Fass, Gal Zizelski Valenci, Mor Rubinstein, Efrat Rorman, Israel Nissan, Efrat Glick-Saar, Omri Nayshool, Gideon Rechavi, Ella Mendelson, Orna Mor |
| EPI_ISL_915876, EPI_ISL_916085 | Lighthouse Lab in Alderley Park | Wellcome Sanger Institute for the COVID-19 Genomics UK (COG-UK) Consortium | Jacquelyn Wynn, Mairead Hyland, The Lighthouse Lab in Alderley Park and Alex Alderton, Roberto Amato, Sonia Goncalves, Ewan Harrison, David K. Jackson, Ian Johnston, Dominic Kwiatkowski, Cordelia Langford, John Sillitoe on behalf of the Wellcome Sanger Institute COVID-19 Surveillance Team |
| EPI_ISL_927125 | Department of Virus and Microbiological Special Diagnostics, Statens Serum Institut, Copenhagen, Denmark | Aalborg University | Danish Covid-19 Genome Consortium |
| EPI_ISL_939446 | Lighthouse Lab in Glasgow | Wellcome Sanger Institute for the COVID-19 Genomics UK (COG-UK) Consortium | Harper VanSteenhouse, Yumi Kasai, David Gray, Carol Clugston, Anna Dominiczak and Alex Alderton, Roberto Amato, Sonia Goncalves, Ewan Harrison, David K. Jackson, Ian Johnston, Dominic Kwiatkowski, Cordelia Langford, John Sillitoe on behalf of the Wellcome Sanger Institute COVID-19 Surveillance Team |
| EPI_ISL_972129 | Department of Virus and Microbiological Special Diagnostics, Statens Serum Institut, Copenhagen, Denmark | Aalborg University | Danish Covid-19 Genome Consortium |
