## Supplemental Table 2 for "CoVizu: Rapid analysis and visualization of the global diversity of SARS-CoV-2 genomes"

We gratefully acknowledge the following Authors from the Originating laboratories responsible for obtaining the specimens, as well as the Submitting laboratories where the genome data were generated and shared via GISAID, on which this research is based.

All Submitters of data may be contacted directly via www.gisaid.org

Authors are sorted alphabetically.

| Accession ID | Originating Laboratory | Submitting Laboratory | Authors |
|---|---|---|---|
| EPI_ISL_1034801, EPI_ISL_1034913 | Landesamt für Verbraucherschutz Sachsen Anhalt, Magdeburg | Institute of Medical Microbiology and Hospital Hygiene | Prof. Dr. Achim Kaasch, Aljoscha Tersteegen |
| EPI_ISL_1039136 | HELIX LLC | WHO National Influenza Centre Russian Federation | Andrey Komissarov, Artem Fadeev, Anna Ivanova, Kseniya Komissarova, Dmitry Bazhenov, Tamila Musaeva, Maria Timofeeva, Veronika Eder, Maria Pisareva, Daria Danilenko, Ksenia Safina, Elena Nabieva, Georgii Bazykin, Dmitry Lioznov |
| EPI_ISL_1092265 | Protzer Lab | Protzer Lab, Laboratory for Functional Genome Analysis, Dept. Genomics, Gene Center of the LMU Munich | Ulrike Protzer, Dieter Hoffmann, Till Bunse, Eva C.Schulte, Elisabeth Esser, Stefan Krebs, Alexander Graf, Helmut Blum |
| EPI_ISL_1149356 | amedes MVZ Halle/Leipzig | Robert Koch Institute | unknown |
| EPI_ISL_1149433, EPI_ISL_1149464 | amedes Göttingen | Robert Koch Institute | unknown |
| EPI_ISL_1154206 | Praxisgemeinschaft für Laboratoriumsmedizin Labor Blumenstraße; Praxis Dr. Sel und Dr. Späte | Robert Koch Institute | unknown |
| EPI_ISL_1156917 | amedes MVZ Halle/Leipzig | Robert Koch Institute | unknown |
| EPI_ISL_1255155 | West African Centre for Cell Biology of Infectious Pathogens (WACCBIP), University of Ghana, Accra, Ghana | West African Centre for Cell Biology of Infectious Pathogens (WACCBIP), University of Ghana, Volta Road, Legon-Accra, Ghana | Collins M. Morang'a, Joyce M. Ngoi, Evelyn B. Quansah, Samirah Saiid, Dominic S.Y. Amuzu, Vincent Appiah, Philip M. Soglo, Vanessa Magnussen, Aisha Mohammed, Kesego Tapela, Nelson Kibinge, Abdoulaye B Diallo, Frederick Kumi-Ansah, Theophilus Odoom, Oliver D Boakye5, Emmanuella Amoako4, Abdul-Karim Abass, , Samuel Kaba Akoriyea, Frederick Tei-Maya, Lucas N. Amenga-Etego, Dam Kenneth Mibut, Yaw Bediako, Benjamin Demah Nuertey, Gordon A Awandare, Peter K Quashie, Gordon A Awandare, Yaw Bediako |
| EPI_ISL_1279884 | Landesamt für Verbraucherschutz Sachsen Anhalt, Magdeburg | Institute of Medical Microbiology and Hospital Hygiene | Prof. Dr. Achim Kaasch, Aljoscha Tersteegen |
| EPI_ISL_1301394 | MSHS Clinical Microbiology Laboratories | MSHS Pathogen Surveillance Program | Ana S. Gonzalez-Reiche, Hala Alshammary, Mitchell J. Sullivan, Brianne Ciferri, Ajay Obla, Angela Amoako, Mahmoud Awawda, Daniel Floda, Julia Matthews, Ashley Salimbangon, Levy Sominsky, Katherine Beach, Kayla Russo, Charles Gleason, Shelcie Fabre, Giulio Kleiner, Zenab Khan, Bremy Alburquerque, Adriana van de Guchte, Komal Srivastava, Matthew M. Hernandez, Jayeeta Dutta, Denise Jurczyszak, Nancy Francoeur, Betsaida Salom Melo, Irina Oussenko, Gintaras Deikus, Juan Soto, Shwetha Hara Sridhar, Ying-Chih Wang, Kathryn Twyman, Deena R. Altman, Robert Sebra, Adolfo García-Sastre, Marta Luksza, Gopi Patel, Sarah Schaefer, Melissa Gitman, Michael D. Nowak, Alberto Paniz-Mondolfi, Emilia Mia Sordillo, Viviana Simon, Harm van Bakel |
| EPI_ISL_1352210 | SYNLAB Jena Oncoscreen | Robert Koch Institute | unknown |
| EPI_ISL_1361757 | Landesamt für Verbraucherschutz Sachsen Anhalt, Magdeburg | Institute of Medical Microbiology and Hospital Hygiene | Prof. Dr. Achim Kaasch, Aljoscha Tersteegen |
| EPI_ISL_1380311 | CSIR-Centre for Cellular and Molecular Biology | CSIR-Centre for Cellular and Molecular Biology-INSACOG | Payel Mukherjee,Pratheusa Maccha,Namami Gaur,Lamuk Zaveri,Tulasi Nagabandi,Purushotham Vodnala,Blessy B John,Viswagithe S L,B Himasri,Sofia Banu,Priya Singh,Archana Bharadwaj Siva,Karthik Bharadwaj Tallapaka,Rakesh K Mishra,Divya Tej Sowpati |
| EPI_ISL_1436787 | amedes MVZ Jena | Robert Koch Institute | unknown |
| EPI_ISL_1437126 | SYNLAB Jena Oncoscreen | Robert Koch Institute | unknown |
| EPI_ISL_1438720, EPI_ISL_1438825 | LabKom - Labor an der Salzbrücke MVZ GmbH | Robert Koch Institute | unknown |
| EPI_ISL_1919433 | HELIX LLC | WHO National Influenza Centre Russian Federation | Andrey Komissarov, Artem Fadeev, Kseniya Komissarova, Oula Masour, Kirill Varchenko, Mikhail Bakaev, Tamila Musaeva, Maria Timofeeva, Veronika Eder, Maria Pisareva, Nikita Yolshin, Daria Danilenko, Ksenia Safina, Elena Nabieva, Georgii Bazykin, Dmitry Lioznov |
| EPI_ISL_2038866 | Federal Budget Health Care Institution "Center of Hygiene and Epidemiology in Tambov region" | Group of Genomics and Postgenomic Technologies of Central Research Institute of Epidemiology | Samoilov AE, Kaptelova VV, Korneenko EV, Saenko SS, Smirnova YS, Nadtoka MI, Belova MV, Speranskaya AS, Tivanova EV, Kondrasheva LY, Akimkin VG |
| EPI_ISL_2156427 | Cagayan Valley Medical Center Molecular Laboratory | Philippine Genome Center | Francis A. Tablizo, Kenneth M. Kim, Carlo M. Lapid, Marc Jerrone R. Castro, Maria Sofia L. Yangzon, Elcid Aaron R. Pangilinan, Benedict A. Maralit, Marc Edsel C. Ayes, Eva Maria Cutiongco-de la Paz, Alethea R. de Guzman, Jan Michael C. Yap, Jo-Hannah S. Llames, Sheila Mae M. Araiza, Kris P. Punayan, Irish Coleen A. Asin, Candice Francheska B. Tambaoan, Asia Louisa U. Chong, Karol Sophia Agape R. Padilla, Rianna Patricia S. Cruz, El King D. Morado, Joshua Gregor A. Dizon, Razel Nikka M. Hao, Arianne A. Zamora, Devon Ray Pacial, Juan Antonio R. Magalang, Marissa Alejandria, Celia Carlos, Anna Ong-Lim, Edsel Maurice Salvaña, John Q. Wong, Jaime C. Montoya, Maria Rosario Singh-Vergeire and Cynthia P. Saloma |
| EPI_ISL_2213652, EPI_ISL_2223184, EPI_ISL_2224086 | Houston Methodist Hospital | Houston Methodist Hospital | Randall J. Olsen, Paul A. Christensen, S. Wesley Long, Sishir Subedi, Robert Olson, Marcus Nguyen, James J. Davis, Matthew Ojeda Saavedra, Prasanti Yerramilli, Layne Pruitt, Kristina Reppond, Madison N. Shyer, Jessica Cambric, Ryan Gadd, Ilya J. Finkelstein, Jimmy Gollihar, and James M. Musser |
| EPI_ISL_2433770 | National Institute for Communicable Diseases of the National Health Laboratory Service | National Institute for Communicable Diseases of the National Health Laboratory Service | Amoako DG, Scheepers C, Mohale T, Ntuli N, Mahlangu B, Ismail A, Bhiman JN |
| EPI_ISL_2433790 | LANCET, LABORATORIES | National Institute for Communicable Diseases of the National Health Laboratory Service | Amoako DG, Scheepers C, Mohale T, Ntuli N, Mahlangu B, Ismail A, Bhiman JN |
| EPI_ISL_2433797 | National Institute for Communicable Diseases of the National Health Laboratory Service | National Institute for Communicable Diseases of the National Health Laboratory Service | Amoako DG, Scheepers C, Mohale T, Ntuli N, Mahlangu B, Ismail A, Bhiman JN |
| EPI_ISL_2433850, EPI_ISL_2433854, EPI_ISL_2433872, EPI_ISL_2433889 | LANCET, LABORATORIES | National Institute for Communicable Diseases of the National Health Laboratory Service | Amoako DG, Scheepers C, Mohale T, Ntuli N, Mahlangu B, Ismail A, Bhiman JN |
| EPI_ISL_2444064 | National Institute for Communicable Diseases of the National Health Laboratory Service | National Institute for Communicable Diseases of the National Health Laboratory Service | Amoako DG, Scheepers C, Mohale T, Ntuli N, Mahlangu B, Ismail A, Bhiman JN |
| EPI_ISL_2617216 | National Health Laboratory Service, South Africa | KRISP, KZn Research Innovation and Sequencing Platform | Giandhari Jennifer, Pillay Sureshnee, Yajna Ramphal, Naidoo Yeshnee, Tshabuila Derek,Tegally Houriiyah, San James, Wilkinson Eduan, de Oliveira Tulio |
| EPI_ISL_467499, EPI_ISL_482716, EPI_ISL_482717, EPI_ISL_482718, EPI_ISL_482851, EPI_ISL_482852, EPI_ISL_482855, EPI_ISL_482857, EPI_ISL_482859 | Molecular Diagnostics Services (MDS) | KRISP, KZN Research Innovation and Sequencing Platform | Giandhari J, Pillay S, Lessells R, Chimukangara B, Mdlalose K, York D, Khan S, Tegally H, Wilkinson E, de Oliveira T |
| EPI_ISL_498055 | NHLS-IALCH | KRISP, KZN Research Innovation and Sequencing Platform | Giandhari J, Pillay S, Lessells R, Chimukangara B, Mdlalose K, York D, Khan S, Tegally H, Wilkinson E, de Oliveira T |
| EPI_ISL_500460 | Laboratório Central de Saúde Pública do Estado de Pernambuco (LACEN-PE) | WallauLab, Aggeu Magalhaes Institute | Marcelo Henrique Santos Paiva, Duschinka Ribeiro Duarte Guedes, Cássia Docena, Matheus Filgueira Bezerra, Filipe Zimmer Dezordi, Laís Ceschini Machado, Larissa Krokovsky, Elisama Helvecio, Alexandre Freitas da Silva, Luydson Richardson Silva Vasconcelos, Antonio Mauro Rezende, Severino |

| | | | |
|---|---|---|---|
| | | | Jefferson Ribeiro da Silva, Kamila Gaudêncio da Silva Sales, Bruna Santos Lima Figueiredo de Sá, Derciliano Lopes da Cruz, Claudio Eduardo Cavalcanti, Armando de Menezes Neto, Caroline Targino Alves da Silva, Renata Pessôa Germano Mendes, Maria Almerice Lopes da Silva, Tiago Gräf, Paola Cristina Resende, Gonzalo Bello, Michelle da Silva Barros, Wheverton Ricardo Correia do Nascimento, Rodrigo Moraes Loyo Arcoverde, Luciane Caroline Albuquerque Bezerra, Sinval Pinto Brandão Filho, Constância Flávia Junqueira Ayres, Gabriel Luz Wallau on behalf of the Fiocruz COVID-19 Genomic Surveillance Network |
| EPI_ISL_509236, EPI_ISL_509330, EPI_ISL_509360, EPI_ISL_509368 | NHLS-IALCH | KRISP, KZN Research Innovation and Sequencing Platform | Giandhari J, Pillay S, Lessells R, Mdlalose K, York D, Tegally H, Wilkinson E, de Oliveira T |
| EPI_ISL_514417, EPI_ISL_514421, EPI_ISL_515152, EPI_ISL_515159, EPI_ISL_515179 | National Institute for Communicable Diseases of the National Health Laboratory Service | National Institute for Communicable Diseases of the National Health Laboratory Service | Allam M, Ismail A, Khumalo Z, Kwenda S, Mtshali P, Mnyameni F, Mohale T, Bhiman JN |
| EPI_ISL_515607, EPI_ISL_515611, EPI_ISL_515639, EPI_ISL_515757, EPI_ISL_515767, EPI_ISL_515774, EPI_ISL_515784 | NHLS-IALCH | KRISP, KZN Research Innovation and Sequencing Platform | Giandhari J, Pillay S, Lessells R, Mdlalose K, York D, Khan S, Tegally H, Wilkinson E, de Oliveira T |
| EPI_ISL_515863 | Medical Disagnostics Services (MDS) | KRISP, KZN Research Innovation and Sequencing Platform | Giandhari J, Pillay S, Lessells R, ChimukangaraB, Mdlalose K, York D, Khan S, Tegally H, Wilkinson E, de Oliveira T |
| EPI_ISL_529732, EPI_ISL_529788, EPI_ISL_535395, EPI_ISL_535444, EPI_ISL_535447, EPI_ISL_535461, EPI_ISL_535572 | NHLS-IALCH | KRISP, KZN Research Innovation and Sequencing Platform | Giandhari J, Pillay S, Lessells R, Mdlalose K, York D, Khan S, Tegally H, Wilkinson E, de Oliveira T |
| EPI_ISL_596257 | HELIX LCC | WHO National Influenza Centre Russian Federation | Andrey Komissarov, Artem Fadeev, Anna Ivanova, Kseniya Komissarova, Dmitry Bazhenov, Daria Danilenko |
| EPI_ISL_602622 | AHRI-Sigal | KRISP, KZN Research Innovation and Sequencing Platform | Gazy I, Sigla, Karim F, Cele S, Giandhari J, Pillay S, Tegally H, Wilkinson E, de Oliveira T |
| EPI_ISL_602624, EPI_ISL_602625 | AHRI-Sigal | KRISP, KZN Research Innovation and Sequencing Platform | Gazy I, Sigl A, Karim F, Cele S, Giandhari J, Pillay S, Tegally H, Wilkinson E, de Oliveira T |
| EPI_ISL_602659, EPI_ISL_602668, EPI_ISL_602669, EPI_ISL_602919 | NHLS-IALCH | KRISP, KZN Research Innovation and Sequencing Platform | Giandhari J, Pillay S, Lessells R, Mdlalose K, York D, Khan S, Tegally H, Wilkinson E, de Oliveira T |
| EPI_ISL_622918 | National Institute for Communicable Diseases of the National Health Laboratory Service | National Institute for Communicable Diseases of the National Health Laboratory Service | Allam M, Ismail A, Khumalo Z, Kwenda S, Mtshali P, Mnyameni F, Mohale T, Subramoney K, Bhiman JN |
| EPI_ISL_622953, EPI_ISL_622957 | Lancet Laboratories | National Institute for Communicable Diseases of the National Health Laboratory Service | Allam M, Ismail A, Khumalo Z, Kwenda S, Mtshali P, Mnyameni F, Mohale T, Subramoney K, Bhiman JN |
| EPI_ISL_634989, EPI_ISL_635006 | National Health Laboratory Service - Inkosi Albert Luthuli Central Hospital (NHLS-IALCH) | KRISP, KZN Research Innovation and Sequencing Platform | Giandhari J, Pillay S, Lessells R, Mdlalose K, York D, Khan S, Tegally H, Wilkinson E, de Oliveira T |
| EPI_ISL_660173 | NHLS-IALCH | KRISP, KZN Research Innovation and Sequencing Platform | Gazy I, Sigal A, Karim F, Cele S, Giandhari J, Pillay S, Tegally H, Wilkinson E, de Oliveira T |
| EPI_ISL_667687 | Pathogen Genomics Center, National Institute of Infectious Diseases | Pathogen Genomics Center, National Institute of Infectious Diseases | Tsuyoshi Sekizuka, Kentaro Itokawa, Rina Tanaka, Masanori Hashino, Makoto Kuroda |
| EPI_ISL_700416 | Zoar Clinic wc ZOA | NHLS/UCT | Arash Iranzadeh, Deelan Doolabh, Lynn Tyers, Bruna Galvao, Innocent Mudau, Marvin Hsiao, Kruger Marais, Diana Hardie, Stephen Korsman, Carolyn Williamson |
| EPI_ISL_700495 | Kwanokuthula CDC wc KWA | NHLS/UCT | Arash Iranzadeh, Deelan Doolabh, Lynn Tyers, Bruna Galvao, Innocent Mudau, Marvin Hsiao, Kruger Marais, Diana Hardie, Stephen Korsman, Carolyn Williamson |
| EPI_ISL_700528 | Plettenberg Bay Clinic wc PLC | NHLS/UCT | Arash Iranzadeh, Deelan Doolabh, Lynn Tyers, Bruna Galvao, Innocent Mudau, Marvin Hsiao, Kruger Marais, Diana Hardie, Stephen Korsman, Carolyn Williamson |
| EPI_ISL_700537 | Kwanokuthula CDC wc KWA | NHLS/UCT | Arash Iranzadeh, Deelan Doolabh, Lynn Tyers, Bruna Galvao, Innocent Mudau, Marvin Hsiao, Kruger Marais, Diana Hardie, Stephen Korsman, Carolyn Williamson |
| EPI_ISL_792309 | Laboratorio del Hospital El Cruce Dr. Néstor C. Kirchner | Área de Secuenciación del Laboratorio de Virología del Hospital de Niños Dr. Ricardo Gutierrez on behalf of 'Proyecto Argentino Interinstitucional de genomica de SARS-CoV-2' (PAIS Consortium) | Nabaes Jodar, MS; Goya, S; Natale, MI; Lusso, S; Zubieta, M; Rahhal, M; Valinotto, LE; Viegas, M. |
| EPI_ISL_792394 | Plataforma de Servicios Biotecnólogicos: UTTIPP/PSB , Universidad Nacional de Quilmes. | Área de Secuenciación del Laboratorio de Virología del Hospital de Niños Dr. Ricardo Gutierrez on behalf of 'Proyecto Argentino Interinstitucional de genomica de SARS-CoV-2' (PAIS Consortium) | Nabaes Jodar, MS; Goya, S; Natale, MI; Lusso, S; Farina, H; Goñi, S; Cardama, G; Castello, A; Valinotto, LE; Viegas, M. |
| EPI_ISL_831294 | Texas Department of State Health Services | Texas Department of State Health Services | Anita Pokharel, Bonnie Oh, James Daniel Bonser, Rashmi Tuladhar, Mayela Pedrueza, Jenny Zhang, Maliha Rahman, Myong Koag, Chung Wang, Rachel Lee, Grace Kubin |
| EPI_ISL_872936 | WHO National Influenza Centre Russian Federation | WHO National Influenza Centre Russian Federation | Andrey Komissarov, Artem Fadeev, Anna Ivanova, Kseniya Komissarova, Dmitry Bazhenov, Mikhail Bakaev, Daria Danilenko, Ksenia Safina, Elena Nabieva, Georgii Bazykin, Dmitry Lioznov |
| EPI_ISL_978238 | Texas Department of State Health Services | Texas Department of State Health Services | Bonnie Oh, Anita Pokharel, James Daniel Bonser, Myong Koag, Chung Wang, Rachel Lee, Grace Kubin, Rashmi Tuladhar, Mayela Pedrueza, Maliha Rahman, Jenny Zhang |
